# Human CD29+/CD56+ myogenic progenitors display tenogenic differentiation potential and facilitate tendon regeneration

**DOI:** 10.1101/2024.04.29.591676

**Authors:** Xiexiang Shao, Xingzuan Lin, Hao Zhou, Minhui Wang, Lili Han, Xin Fu, Sheng Li, Siyuan Zhu, Shenao Zhou, Wenjun Yang, Jianhua Wang, Ping Hu

## Abstract

Tendon injury occurs at high frequency and is difficult to repair. Identification of human stem cells being able to regenerate tendon will greatly facilitate the development of regenerative medicine for tendon injury. We identified human CD29+/CD56+ myogenic progenitors having tendon differentiation potential both in vitro and in vivo. Transplantation of human CD29+/CD56+ myogenic progenitors contributes to injured tendon repair and thus improves locomotor function. Interestingly, the tendon differentiation potential in mouse muscle stem cells is minimal and the higher TGFβ signaling level may be the key for the distinct feature of human CD29+/CD56+ myogenic progenitors. These findings reveal that human CD29+/CD56+ myogenic progenitors are bi-potential adult stem cells and can serve as a new source for tendon regeneration.

## Introduction

Skeletal muscle is a tissue with great regeneration ability due to the existence of muscle stem cells (MuSCs). MuSCs are adult stem cells located at the periphery of myofibers between the basal lamina and the plasmalemma of the myofibers and responsible for muscle regeneration^1,2^. MuSCs have been considered to be unipotent stem cells to have a sole differentiation potential to myofibers^3^. MuSCs undergo expansion and differentiate to multinuclei myofibers after injury in vivo. MuSCs have remarkable abilities to support muscle regeneration. After transplantation, the engrafted mouse MuSCs go through active expansion and regenerate myofibers^4^. Isolation by FACS and expansion of MuSCs have been reported from several species including mouse, pig, and human^5,6^. Single cell sequencing analysis from human skeletal muscles have also revealed the existence of Pax7+ myogenic progenitors ^6,7^. Due to the different motion patterns, the regeneration capacity may be different between human and rodents. Current investigations have suggested that human myogenic progenitor cells didn’t share same markers as that in mice^8^, and the expression pattern of oxidative enzymes and cytokines between these two species are also different^9^, suggesting that human muscle progenitor cells may have distinct features from mice.

FACS method to isolate cells with myogenic differentiation potential from human muscle biopsies has been established. It has been reported that CD56 (NCAM)+ and CD56+ CD29+ cells from skeletal muscle display myogenic differentiation potential^9,10,11,12^. The CD56+ CD34+ progenitor cells isolated from human skeletal muscle biopsies have been reported to have chondrogenic, adipogenic, and osteogenic potentials besides myogenic potential^13–15^. CD56+CD34-progenitor cells are free of adipogenic potential^11^. The differentiation potential of human myogenic progenitors remains to be further explored.

Skeletal muscle directly connects to tendons which is responsible for transmitting forces from skeletal muscle to bone to generate active movement. Tendinopathy affects more than 10% of the population under 45 and compromises the tendon functions^16,17^. Tendon injury healing is slow and incomplete due to the low number of cells in tendon and the hypovascular and anaerobic environment^18,19^. Tendon stem/progenitor cells (TDSCs) is a cell population derived from tendon and considered to be a subgroup of mesenchymal stem cells that have abilities to improve tendon injury healing^20,21^. However, the number of TDSCs in tendon is low and retrieving TDSCs is invasive and exacerbates tendon injury. Morphology and proliferation ability loss in the culture system also hampers the efforts to obtain sufficient amounts of active TDSCs. To find more cell types supporting tendon regeneration will facilitate the development of regenerative medicine to treat tendon injury. The activity of tenogenesis is tightly regulated by many signaling pathways, especially for TGFβ signaling^22^. TGFβ signaling is indispensable for tendon development^23^, and it could systematically promote the tenogenic differentiation of stem cells^24,25^. The downstream effectors SMAD2 and SMAD3 are able to activate the transcription of tendon-specific genes which further facilitates tendon development and tenogenic differentiation^22,26,27^. After tendon injury, TGFβ signaling promotes the proliferation, migration and differentiation of TDSCs^28^, increases tendon collagen synthesis^29^, and contributes to matrix anabolism for tendon remodeling^30^.

Here we found that human CD29+/CD56+ myogenic progenitors can be differentiated to tendon cells both in vitro and in vivo. Transplantation of human CD29+/CD56+ myogenic progenitors to injured tendon in mice improved the tendon regeneration, suggesting its bipotential ability. Interestingly, the tendon differentiation potential in mouse muscle stem cells is minimal and the higher TGFβ signaling level may be the key for the distinct feature of human CD29+/CD56+ myogenic progenitors.

## Results

### The CD29+/CD56+ cells isolated from human muscle biopsies display robust features of myogenic progenitors

To analyze the cell components of human skeletal muscle, single cell sequencing analysis was performed using skeletal muscle biopsy. A total of 57,193 cells were included for analysis. The cells were grouped to 9 cell populations (Fig. 1a and b). Consistent with the previous single cell sequencing results from human muscles^31^, Fibro/Adipogenic progenitors (FAPs), endothelial cells, myocytes, and myogenic progenitors were among the identified cell types (Fig. 1a and b). Especially, myogenic progenitors accounted for 12.4% of all the mononuclear cells, while tenocytes only accounted for 0.06% (Fig. 1c). To investigate the identity of CD29+/CD56+ cells, joint expression analysis was performed (Fig. 1d). The scRNA-seq data revealed that all the CD29+/CD56+ cells were myogenic progenitors, which occupied 19.3% of all the myogenic progenitors (Fig. 1e). However, there existed no tenocytes with CD29+/CD56+ (Fig. 1d). Combined, the scRNA-seq data revealed human CD29+/CD56+ cells were myogenic progenitors.

**Figure 1.**
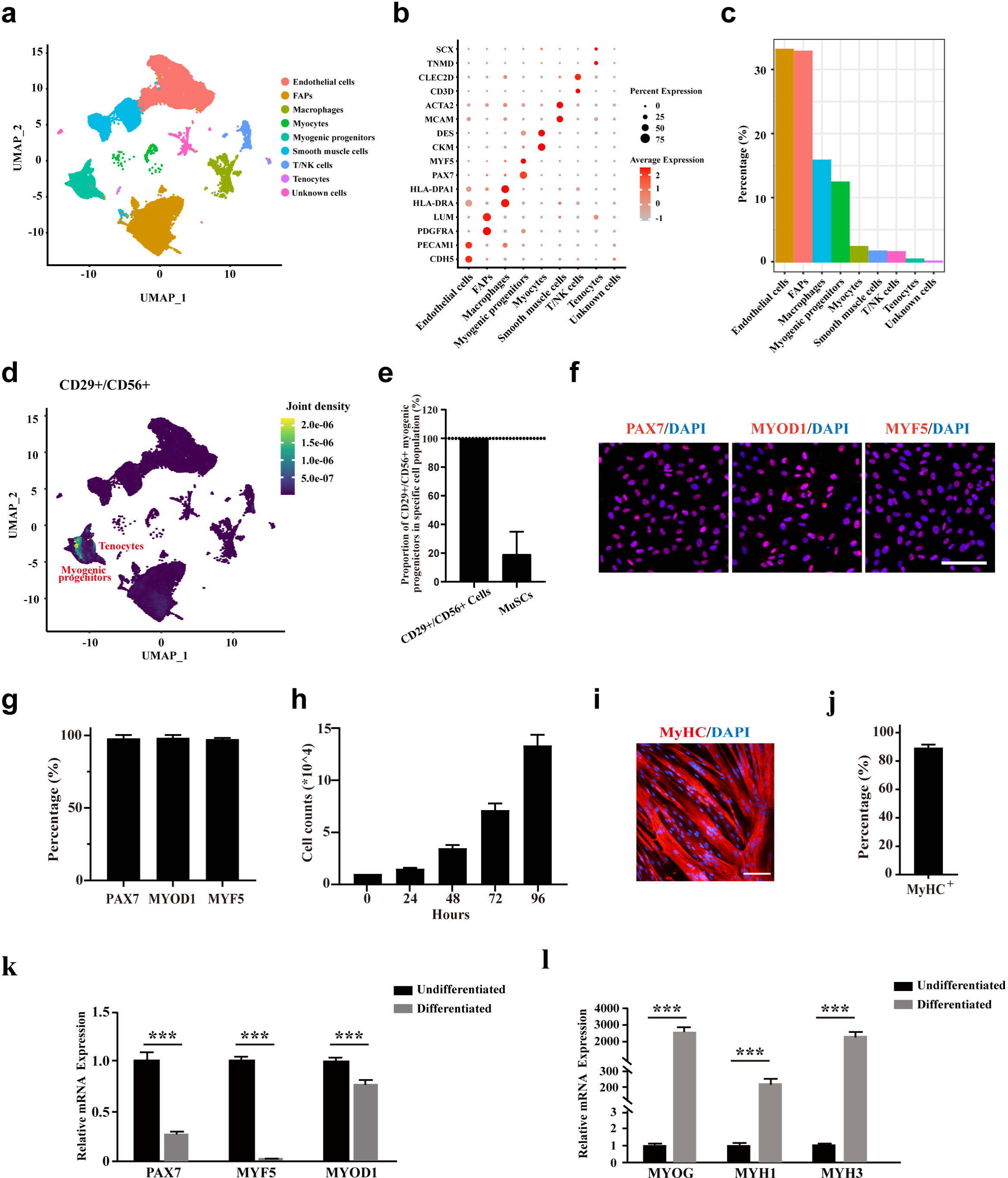
The CD29+/CD56+ cells isolated from human muscle biopsies display robust features of myogenic progenitors. a. UMAP plot of all mononuclear cells isolated from human skeletal muscle which were colored by cell clusters. Three samples with a total of 57,193 cells were included for analysis. b. Dot blot of representative genes in each cell cluster. c. Bar plot of cell proportion for each cell cluster. d. Plot of joint expression analysis for CD29+/CD56+ cells in total mononuclear cells isolated from human skeletal muscle. e. Proportion of CD29+/CD56+ myogenic progenitors in specific cell population. Error bars indicated standard deviation (n=3). f. Immunofluorescence staining of PAX7, MYOD1 and MYF5 in primary human CD29+/CD56+ myogenic progenitors. Scale bars, 100µm. g. Statistical analysis of the percentage of PAX7+, MYOD1+ and MYF5+ cells in the isolated CD29+/CD56+ myogenic progenitors. Error bars indicated standard deviation (n=5). h. Analysis of cell proliferation for human CD29+/CD56+ myogenic progenitors in vitro. 10,000 isolated human CD29+/CD56+ myogenic progenitors were plated for proliferation and counted at each time point. Error bars indicated standard deviation (n=3). i. Immunofluorescence staining of MyHC in myotubes differentiated from human CD29+/CD56+ myogenic progenitors. Primary human CD29+/CD56+ myogenic progenitors were isolated and differentiated to myotubes for 5 days followed by MyHC immunofluorescence staining. Scale bars, 100 µm. j. Statistical analysis of the percentage of nuclei in MyHC+ myotubes after differentiation of the human CD29+/CD56+ myogenic progenitors. Error bars indicated standard deviation (n=5). k. Relative expression level of PAX7, MYF5, and MYOD1 in human CD29+/CD56+ myogenic progenitors (Undifferentiated) and differentiated myotubes (Differentiated). RT-qPCR assays were performed for human CD29+/CD56+ myogenic progenitors before and after myogenic differentiation. GAPDH was served as reference gene. Error bars indicated standard deviation (n=3). *** indicated p<0.001. l. Relative expression level of MYOG, MYH1, and MYH3 in human CD29+/CD56+ myogenic progenitors (Undifferentiated) and differentiated myotubes (Differentiated). RT-qPCR assays were performed for human CD29+/CD56+ myogenic progenitors before and after myogenic differentiation. GAPDH was served as reference gene. Error bars indicated standard deviation (n=3). *** indicated p<0.001.

To confirm the single cell analysis results, we first isolated CD29+/CD56+ myogenic progenitors from human muscle biopsy using FACS as described previously^6,12^. CD31-/CD45-/CD29+/CD56+ cells were collected from the single cell suspension dissociated from muscle tissue. The collected cells were stained with anti-PAX7, anti-MYOD1 and anti-MYF5 antibody to confirm the purity of isolated CD29+/CD56+ cells, and nearly all of the obtained cells were positively stained with these myogenic progenitors markers (Fig. 1f and g). The proliferation assay indicated high in vitro expansion ability of these human CD29+/CD56+ myogenic progenitors (Fig. 1h). When induced to differentiate by 0.4% Ultroser G, human CD29+/CD56+ myogenic progenitors differentiate to myotubes robustly. Approximately 90% of nuclei were present in Myosin heavy chain (MyHC) positive myotubes (Fig. 1i and j). Consistently, the expression of PAX7, MYF5 and MYOD1 were enriched in the myogenic progenitors, and their expression decreased after differentiation as shown by RT-qPCR assays (Fig. 1k). In contrast, the expression of genes marking differentiation such as Myogenin (MYOG), Myosin heavy chain 1 (MYH1), and Myosin heavy chain 3 (MYH3) were significantly up-regulated in differentiated myotubes (Fig. 1l). Taken together, these results suggest that the CD29+/CD56+ cells isolated from human muscle biopsies have robust features of myogenic progenitors.

### Human CD29+/CD56+ myogenic progenitors display tenogenic differentiation potential in vitro

Since previous studies indicated that there could be tenogenic differentiation ability for muscle-derived cells^32–34^, we next explored the tenogenic differentiation potential of CD29+/CD56+ myogenic progenitors. The isolated primary human CD29+/CD56+ myogenic progenitors were induced to differentiate to tenocytes by 100ng/ml GDF5, 100ng/ml GDF7 and 0.2mM ascorbic acid for 12 days. After 12 days of tendon differentiation, the morphology of cells displayed dramatic differences from those undergoing myogenic differentiation (SFig. 1a). Furthermore, the expression of tendon markers such as TNC and SCX was significantly increased (Fig. 2a, b, and c). Moreover, some other tendon related genes, such as COL I, MKX, THBS4, and COMP, were also enriched after tendon differentiation (Fig. 2c). In contrast, the expression of these genes was not enriched upon myogenic differentiation (Fig. 2a, b, and c). The expression of genes marking myogenic differentiation such as MYOG and MyHC were only detected in a small portion of cells after tenogenic differentiation (SFig. 1b and c). Compared to the over 90% of differentiation efficiency upon myogenic differentiation, only about 20% of cells showed MYOG and MyHC expression after 12 days of tenogenic induction (SFig. 1b and c). The genes marking differentiated myotubes such as MYH1, MYH3, DESMIN, and MYL1 showed moderate elevation after tenogenic differentiation, while dramatic upregulation of these genes was observed after myogenic differentiation (SFig. 1d and e). These results combined suggest that human CD29+/CD56+ myogenic progenitors are capable of tendon differentiation in vitro.

**Figure 2.**
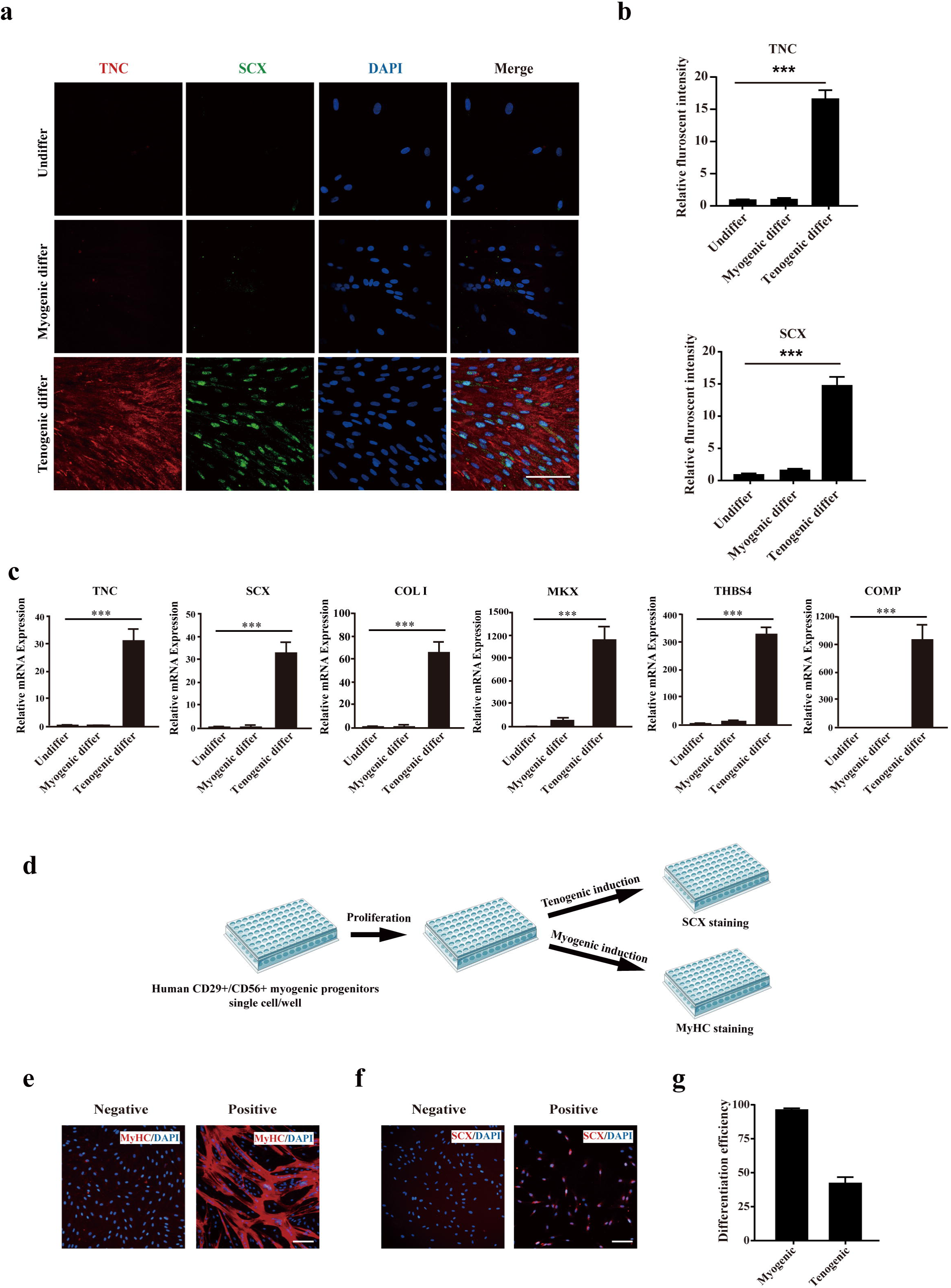
Human CD29+/CD56+ myogenic progenitors display tendon differentiation ability in vitro. a. Immunofluorescence staining of tendon marker TNC and SCX in human CD29+/CD56+ myogenic progenitors induced for myogenic and tenogenic differentiation, respectively. Scale bars, 100 µm. b. Quantification of TNC and SCX fluorescent intensity in human CD29+/CD56+ myogenic progenitors undergone myogenic and tenogenic differentiation, respectively. Error bars indicated standard deviation (n=5). c. Relative expression levels of genes enriched in tendon cells. RT-qPCR assays were performed with human CD29+/CD56+ myogenic progenitors upon myogenic and tenogenic differentiation, respectively. GAPDH was served as reference gene. Error bars indicated standard deviation (n=3). *** indicated p<0.001. d. Scheme of clonal proliferation and differentiation assay. e. Representative immunofluorescence staining images of MyHC as the marker for successful myogenic differentiation. Scale bars, 100 µm. f. Representative immunofluorescence staining images of SCX as the marker for successful tenogenic differentiation. Scale bars, 100 µm. g. Statistical analysis of the myogenic and tenogenic differentiation efficiency of human CD29+/CD56+ myogenic progenitors. Error bars indicated standard deviation (n=3).

To further confirm the results, we next performed clonal analysis. The freshly isolated primary human CD29+/CD56+ myogenic progenitors were seeded to 96 well plate with the concentration of 1 cell/well. The wells of single cell were allowed to proliferate for 10 days followed by tenogenic or myogenic induction. The plates with myogenic induction were differentiated for 4 days and immunofluorescence staining of MyHC was performed to determine myogenic differentiation. The plates with tenogenic induction were differentiated for 12 days and immunofluorescence staining of SCX was performed to determine tenogenic differentiation (Fig. 2d). The number of wells showing positive MyHC staining was counted and the myogenic differentiation efficiency was calculated (Fig. 2e and g). MyHC+ myotubes was observed in over 95% of wells with alive cells (Fig. 2g). Similarly, the number of wells displaying positive SCX staining were counted and the tenogenic differentiation efficiency was calculated (Fig. 2f and g). Approximately 40% of CD29+/CD56+ myogenic progenitors displayed tenogenic differentiation ability (Fig. 2g), suggesting that human CD29+/CD56+ myogenic progenitors have tenogenic differentiation potential. Taken together, these results suggest that human CD29+/CD56+ myogenic progenitors have dual differentiation potentials towards muscle or tendon in vitro.

### Tenocytes differentiated from human CD29+/CD56+ myogenic progenitors display similar expression profile to primary tenocytes

RNA sequencing was then performed to further determine the lineage of the tenocytes differentiated from human CD29+/CD56+ myogenic progenitors. Human CD29+/CD56+ myogenic progenitors were induced towards myogenic or tenogenic differentiation respectively. We also isolated primary human tenocytes as described previously^35^. These cells were subjected for RNA sequencing analysis. The differentiated myotubes and tenocytes displayed distinct expression patterns (Fig. 3a). The differentiated tendon cells share high similarity to fresh isolated tenocytes from human tendons (Fig. 3a), suggesting that human CD29+/CD56+ myogenic progenitors are capable of tendon differentiation. Consistently, two distinct sets of genes representing myogenic markers and tenogenic markers were up-regulated after myogenic induction and tenogenic induction, respectively (Fig. 3b and c). These results suggest that human CD29+/CD56+ myogenic progenitors are capable of dual direction differentiation.

**Figure 3.**
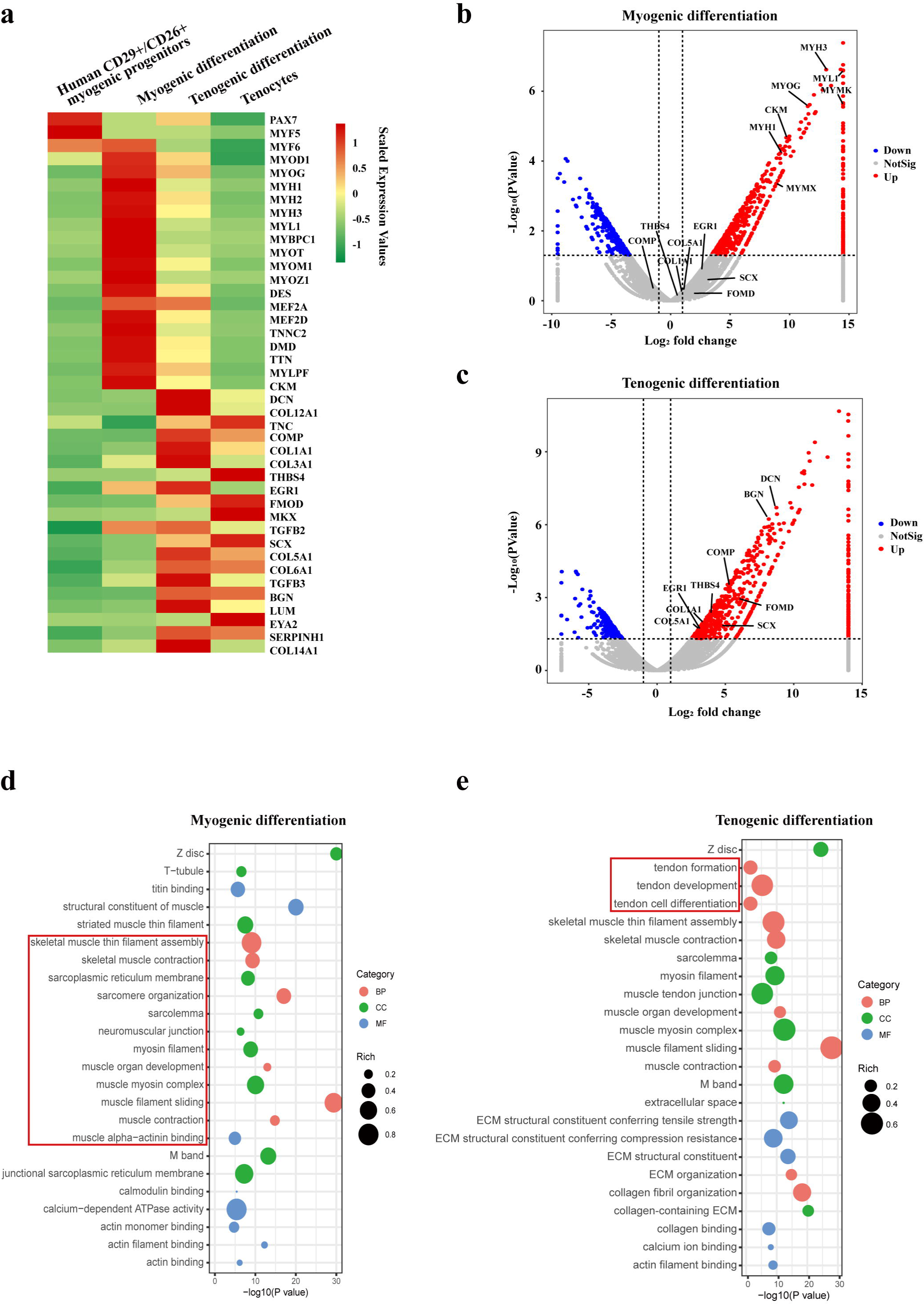
Human CD29+/CD56+ myogenic progenitors have tendon differentiation potential. a. Heat map of gene expression profiles of human CD29+/CD56+ myogenic progenitors, human CD29+/CD56+ myogenic progenitors after myogenic differentiation, human CD29+/CD56+ myogenic progenitors after tenogenic differentiation, and primary tenocytes. b. Volcano plot of genes enriched in myogenic differentiation of human CD29+/CD56+ myogenic progenitors. c. Volcano plot of genes enriched in tenogenic differentiation of human CD29+/CD56+ myogenic progenitors. d. Bubble chart of GO analysis of cellular process up-regulated in myogenic differentiation of human CD29+/CD56+ myogenic progenitors. e. Bubble chart of GO analysis of cellular process up-regulated in tenogenic differentiation of human CD29+/CD56+ myogenic progenitors.

Further GO analysis also displayed the activation of two distinct sets of cell features. Upon the skeletal muscle differentiation, terms related to skeletal muscle functions such as skeletal muscle thin filament assembly, skeletal muscle contraction, muscle organ development, and sarcomere organization were enriched (Fig. 3d), suggesting the muscular identity of the differentiated cells. In contrast, terms related to tendon formation, tendon development, and tendon cell differentiation were enriched after tenogenic differentiation (Fig. 3e), suggesting that tendon identity is achieved in the differentiated cells. Together, these results suggest that human CD29+/CD56+ myogenic progenitors are capable of both myogenic and tenogenic differentiation in vitro.

### Murine MuSCs display poor tenogenic differentiation ability

We next investigated whether the tenogenic differentiation ability is present in rodent muscle stem cells (MuSCs). Mouse MuSCs were isolated by positive marker of Vcam1 as described previously^6^ and induced for tenogenic differentiation. In sharp contrast to human CD29+/CD56+ myogenic progenitors, murine MuSCs failed to be induced to tendon cells upon the same induction condition as that for human CD29+/CD56+ myogenic progenitors, though the myogenic differentiation is as efficient as the human CD29+/CD56+ myogenic progenitors. After myogenic differentiation, 93.9% of nuclei were present in MyHC+ myotubes (Fig. 4a and b). In sharp contrast, no Scx+ cells were observed after 12 days of induction for tenogenic differentiation (Fig. 4a). Consistent with the immunofluorescence staining results, RT-qPCR results revealed that myogenic differentiation marker genes such as MyoG, Myh1, and Myh3 were significantly up-regulated under both myogenic and tenogenic differentiation conditions (Fig. 4c), suggesting that murine MuSCs predominantly commit myogenic differentiation under induction. Different from human CD29+/CD56+ myogenic progenitors, the expression of genes indicating tendon cell fate such as Scx, Tnc, Col I, Mkx, and Thbs4 did not increase after tenogenic differentiation (Fig. 4d), suggesting the failure to induce tenogenic cell fate from murine MuSCs. Together, these results suggest that murine MuSCs display almost no tenogenic differentiation potential in vitro.

**Figure 4.**
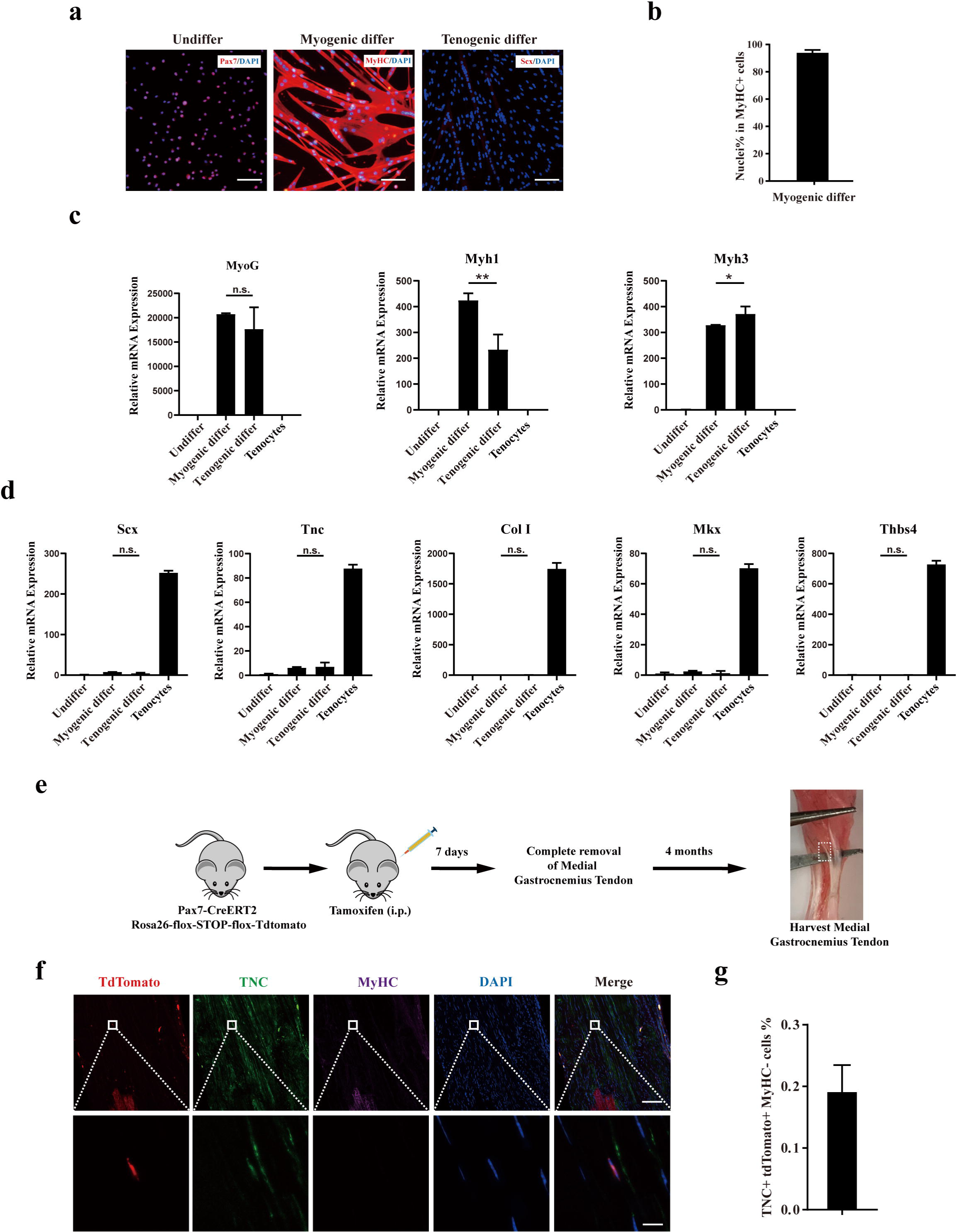
Murine MuSCs display poor tenogenic differentiation ability. a. Immunofluorescence staining of murine MuSCs directed towards myogenic and tenogenic differentiation, respectively. The undifferentiated murine MuSCs were stained with Pax7 and DAPI. Murine MuSCs after myogenic differentiation were stained with MyHC and DAPI. Murine MuSCs after tenogenic differentiation were stained with Scx and DAPI. Scale bars, 100 µm. b. Statistical analysis of the percentage of nuclei in MyHC+ myotubes. Error bars indicated standard deviation (n=5). c. Relative expression levels of myogenic differentiation specific genes MyoG, Myh1, and Myh3. Mouse MuSCs were induced to differentiate towards muscle and tendon, respectively. Mouse MuSCs before and after differentiation together with primary tenocytes were harvested and subjected for RT-qPCR analysis. Gapdh was served as reference gene. Error bars indicated standard deviation (n=3). * indicated p<0.05, ** indicated p<0.01, n.s. indicated n>0.05. d. Relative expression levels of tenogenic differentiation marker genes Scx, Tnc, Col I, Mkx and Thbs4. Mouse MuSCs were induced to differentiate towards muscle and tendon, respectively. Mouse MuSCs before and after differentiation together with primary tenocytes were harvested and subjected for RT-qPCR analysis. Gapdh was served as reference gene. Error bars indicated standard deviation (n=3). n.s. indicated n>0.05. e. Scheme of Pax7+ MuSC progeny lineage tracing assay. f. Immunofluorescence staining of tendon tissue 4 months after injury. tdTomato+ cells indicated the progeny of Pax7+ MuSCs. TNC+ cells indicated tendon cells. MyHC+ cells indicated muscle fibers. DAPI indicated nuclei staining. Merged indicated the merged images of tdTomato, TNC, MyHC, and DAPI. The upper panel indicated the low magnification images. Scale bars, 100 µm. The lower panel indicated the high magnification images of the region label by white square in the upper panel. Scale bars, 10 µm. g. The statistical analysis of the percentage of tdTomato+ TNC+ cell 4 months after tendon injury. Error bars indicated standard deviation (n=5).

We next performed the lineage tracing experiments in mice to further examine the tenogenic potential of mouse MuSCs in vivo (Fig. 4e). Pax7CreERT2 mice were crossed to flox-Stop-flox-tdTomato mice. MuSCs and the descendants of MuSCs will be labeled by tdTomato (SFig. 2a). Tendon injury in mice was generated by mimicking the peroneus longus tendon removal surgery in human (SFig. 2b). In human, it has been reported that the tendon could be regenerated to some extent after the peroneus longus tendon removal surgery based on MRI imaging^32^. In this surgery, injury of the skeletal muscle adjacent to the removed tendon was inevitable. The accompanied skeletal muscle injury could activate MuSCs and make them available for tendon regeneration. To further activate MuSCs to guarantee that sufficient amount of activated MuSCs were available around the tendon injury site, we also injected 15μl of 10μM cardiotoxin (CTX) at the muscle adjacent to the sites where the tendon was removed to induce more muscle injuries and further activated MuSCs. If MuSCs can participate tendon regeneration, tdTomato+ tendon cells would be observed after the repair of tendon injury.

Expectedly, large amount of tdTomato+ myofibers were observed after muscle injury (SFig. 2c-d), suggesting that the tracing system works well. Nevertheless, less than 0.2% tendon cells originated from mouse MuSCs were observed even four months after tendon removal (Fig. 4f-g). These results suggest that murine MuSCs have poor tendon differentiation abilities.

### Transplantation of human CD29+/CD56+ myogenic progenitors facilitates tendon regeneration

We next went on to investigate whether the transplantation of human CD29+/CD56+ myogenic progenitors to mouse can improve tendon regeneration. An approximately 1.5mm long and 0.5mm width transverse incision was performed at 5mm from the calcaneus in Achilles tendon for NOD/SCID immunodeficient mice. Total 50,000 human CD29+/CD56+ myogenic progenitors packed in hydrogel were planted at the incision site (Fig. 5a). As a control, mixture of PBS and hydrogel was transplanted in the SCID recipient mice undergoing the same tendon injury at the cleavage sites. Packing the cells with hydrogel concentrated the transplanted cells at the local injury sites. Two months after transplantation, the tendons carrying transplanted with or without human CD29+/CD56+ myogenic progenitors were harvested, respectively (Fig. 5a).

**Figure 5.**
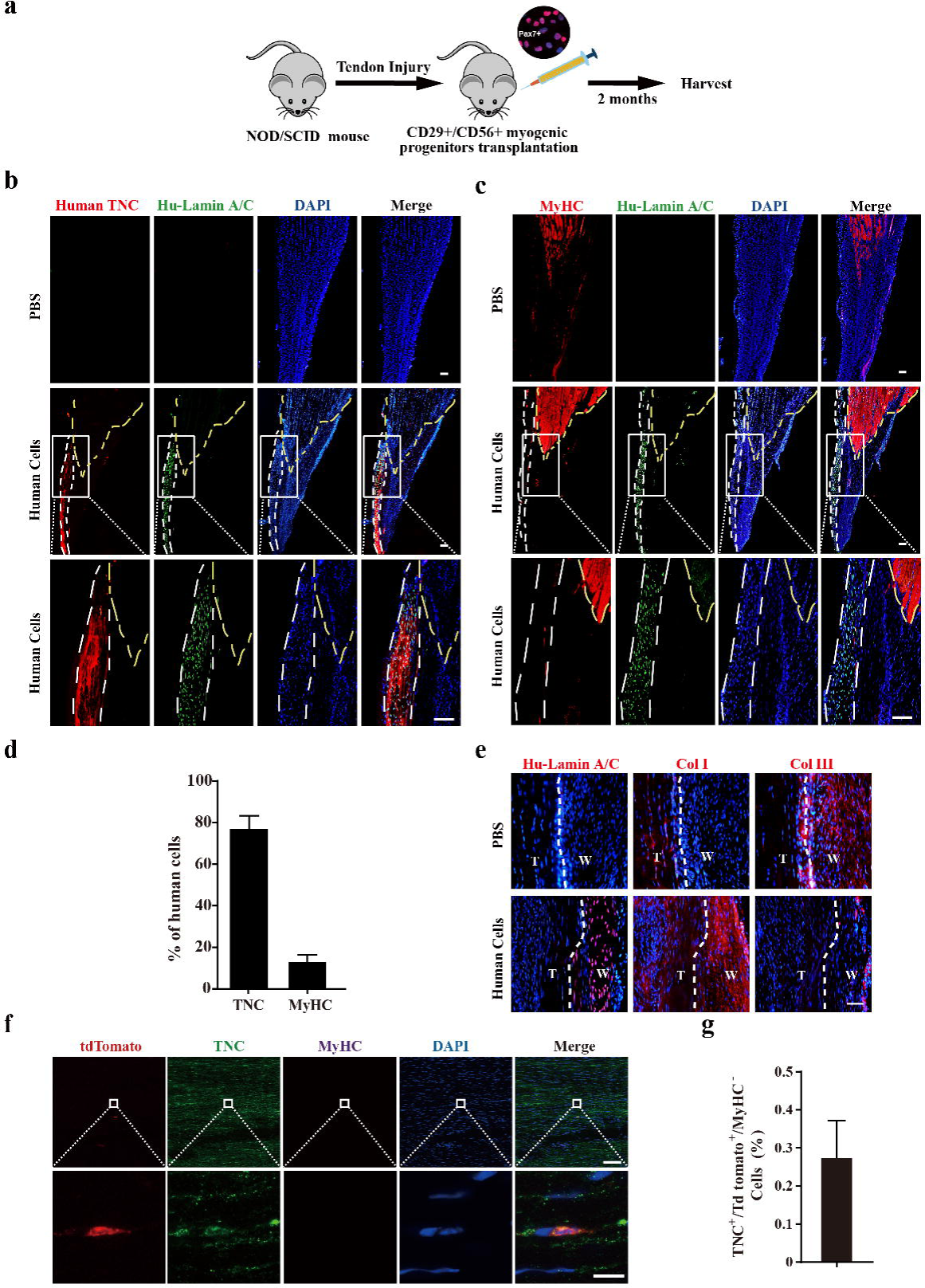
Transplantation of human CD29+/CD56+ myogenic progenitors facilitates tendon regeneration. a. Scheme of transplantation of human CD29+/CD56+ myogenic progenitors for NOD/SCID with tendon injury. b. Immunofluorescence staining of the regenerated tendon-like tissue after transplantation of human CD29+/CD56+ myogenic progenitors. Tendon injury was induced in recipient NOD/SCID mice and 50,000 human CD29+/CD56+ myogenic progenitors were transplanted to the injured tendon at the injured site. The regenerated tendon-like tissue, the connected muscle, and the surrounding soft tissues were harvested to make continuous cryosections. One of the continuous cryosections were subjected for immunofluorescence staining of tendon marker human specific TNC and human specific Lamin A/C. DAPI indicated nuclei staining. Merge indicated the merged images of human TNC, human Lamin A/C, and DAPI. The white lines indicated the location of regenerated tendon-like tissues from human CD29+/CD56+ myogenic progenitors based on human TNC staining. The yellow dashed lines indicated the superimposed location of muscle based on MyHC staining in panel c. Scale bars, 100 µm. c. Immunofluorescence staining of MyHC and human Lamin A/C. The regenerated tendon-like tissue, the connected muscle, and the surrounding soft tissues were harvested and subjected for continuous cryosection. One of the continuous cryosections were stained for MyHC which was specifically expressed in skeletal muscle, and human Lamin A/C to label human cells. DAPI indicated nuclei staining. Merge indicated the merged images of MyHC, human Lamin A/C, and DAPI. The white lines indicated the location of regenerated tendon-like tissue from human CD29+/CD56+ myogenic progenitors based on human TNC staining in panel b. The yellow dashed lines indicated the location of muscle based on MyHC staining. Scale bars, 100µm. d. Statistical analysis of the percentage of human cells expressing skeletal muscle marker MyHC or tendon marker TNC after being transplanted to the injured tendon. Error bars indicated standard deviation (n=5). e. Immunofluorescence staining of Col I, Col III and human Lamin A/C. Two months after human cell transplantation, continuous cryosections containing the regenerated tendon-like tissue and native tendon tissue was stained with Col I, Col III and human Lamin A/C. DAPI indicated the staining of nuclei. T, native tendon tissue; W, wound tendon tissue. Scale bars, 50 µm. f. Immunofluorescence staining of tendon tissue after transplantation of tdTomato+ murine MuSCs. tdTomato indicated the progenies of murine MuSCs. TNC indicated immunofluorescence staining of tendon marker TNC. MyHC indicated immunofluorescence staining of myofiber marker MyHC. DAPI indicated nuclei staining. Merge indicated merged images of tdTomato, TNC, MyHC, and DAPI. The upper panel indicated low magnification images. Scale bars, 100 µm. The lower panel indicated the amplified images of the region indicated by the white square. Scale bars, 10 µm. g. Statistical analysis of TNC+ tdTomato+ cells in tendon tissue after transplantation of murine MuSCs. Error bars indicated standard deviation (n=5).

In mice transplanted with human CD29+/CD56+ myogenic progenitors, continuous cryosections containing muscle and tendon tissues were generated. Immunofluorescence staining was performed to detect tendon and muscle markers with two cryosections adjacent to each other, respectively. The two sets of images obtained on continuous cryosections were superimposed on each other to pinpoint the position of tendons. Immunofluorescence staining of antibody specifically recognizing human TNC indicated the position of regenerated tendon-like tissue from human cells in the harvested tissues (Fig. 5b). The presence of human cells was also illustrated by the immunofluorescence staining with the antibody specifically recognizing human Lamin A/C. In the control PBS injection group, where the mixture of PBS and hydrogel instead human CD29+/CD56+ myogenic progenitors was injected to the incision sites, no human Lamin A/C was detected. These results confirmed the specificity of human TNC and Lamin A/C antibody. Immunofluorescence staining revealed that over 75% of the human cells showed TNC expression (Fig. 5b and d), suggesting that the majority of the transplanted human CD29+/CD56+ myogenic progenitors differentiate to tendon cells in vivo. The immunofluorescence staining of MyHC and human Lamin A/C were performed to detect the muscle cells originated from the transplanted human CD29+/CD56+ myogenic progenitor cells. Only about 12.8% of the human cells detected expressed MyHC (Fig. 5c and d). Moreover, the human cells were predominantly enriched at the tendon region (Fig. 5b). Only a few MyHC+ cells originated from the transplanted human CD29+/CD56+ myogenic progenitors scattered in the muscle region (Fig. 5c). To further confirm the tendon differentiation potential of the transplanted human CD29+/CD56+ myogenic progenitors, immunofluorescence staining of SCX and TNMD was also performed. The majority (80.0% and 74.6%) of transplanted human CD29+/CD56+ myogenic progenitors also expressed SCX and TNMD (SFig. 3a, b and c). Furthermore, Col I was predominantly expressed in the regenerated tendon-like tissue rather than Col III after human cells transplantation (Fig. 5e), indicating human CD29+/CD56+ myogenic progenitors could contribute to structural repair of injured tendon and facilitate the healing process. Taken together, these results suggest that human CD29+/CD56+ myogenic progenitors are capable of tendon differentiation in vivo and contributing to tendon regeneration.

In sharp contrast to human CD29+/CD56+ myogenic progenitors, when 50,000 murine MuSCs constitutively expressing tdTomato were transplanted to pre-injured tendon in NOD/SCID mice under the same condition, less than 0.3% of tdTomato+ TNC+ cells were detected (Fig. 5f and g). However, the myogenic differentiation potential of human CD29+/CD56+ myogenic progenitors and mouse MuSCs was similar. Muscle injury was induced by muscular injection of CTX in tibialis anterior (TA) muscle in NOD/SCID mice. TA muscles were irradiated to kill the local MuSCs as described previously^36^. Transplantation of human CD29+/CD56+ myogenic progenitors and mouse MuSCs to the irradiated pre-injured recipient mice was performed, respectively. TA muscles were harvested after 28 days. Transplantation of both human CD29+/CD56+ myogenic progenitors and murine MuSC displayed robust engraftment efficiency (SFig. 3d-i). These results suggest that human CD29+/CD56+ myogenic progenitors and murine MuSCs have similar myogenic differentiation potential.

Combined, the above results suggest that human CD29+/CD56+ myogenic progenitors have dual differentiation potentials towards myogenesis or tenogenesis in vivo, while murine MuSCs predominantly commit myogenesis.

### Transplantation of human CD29+/CD56+ myogenic progenitors improves locomotor function after tendon injury

We next checked whether transplantation of human CD29+/CD56+ myogenic progenitors could improve tendon functions. First, transmission electron microscope analysis was used to evaluate microstructure of injured tendon two months after transplantation. Larger and denser collagen fibrils of the tendons were identified in transplanted group than control group with PBS injection, although the maturation level of collagen fibrils could still not reach to those in uninjured tendon (Fig. 6a and b). We next examined whether transplantation of human CD29+/CD56+ myogenic progenitors could improve the biomechanical property of tendon. The max failure load and stiffness of the tendons from the transplantation group were significantly better than those from PBS injection control group (Fig. 6c). These results combined suggest that transplantation of human CD29+/CD56+ myogenic progenitors improves the collagen fibril maturation and biomechanical property of the injured tendon.

**Figure 6.**
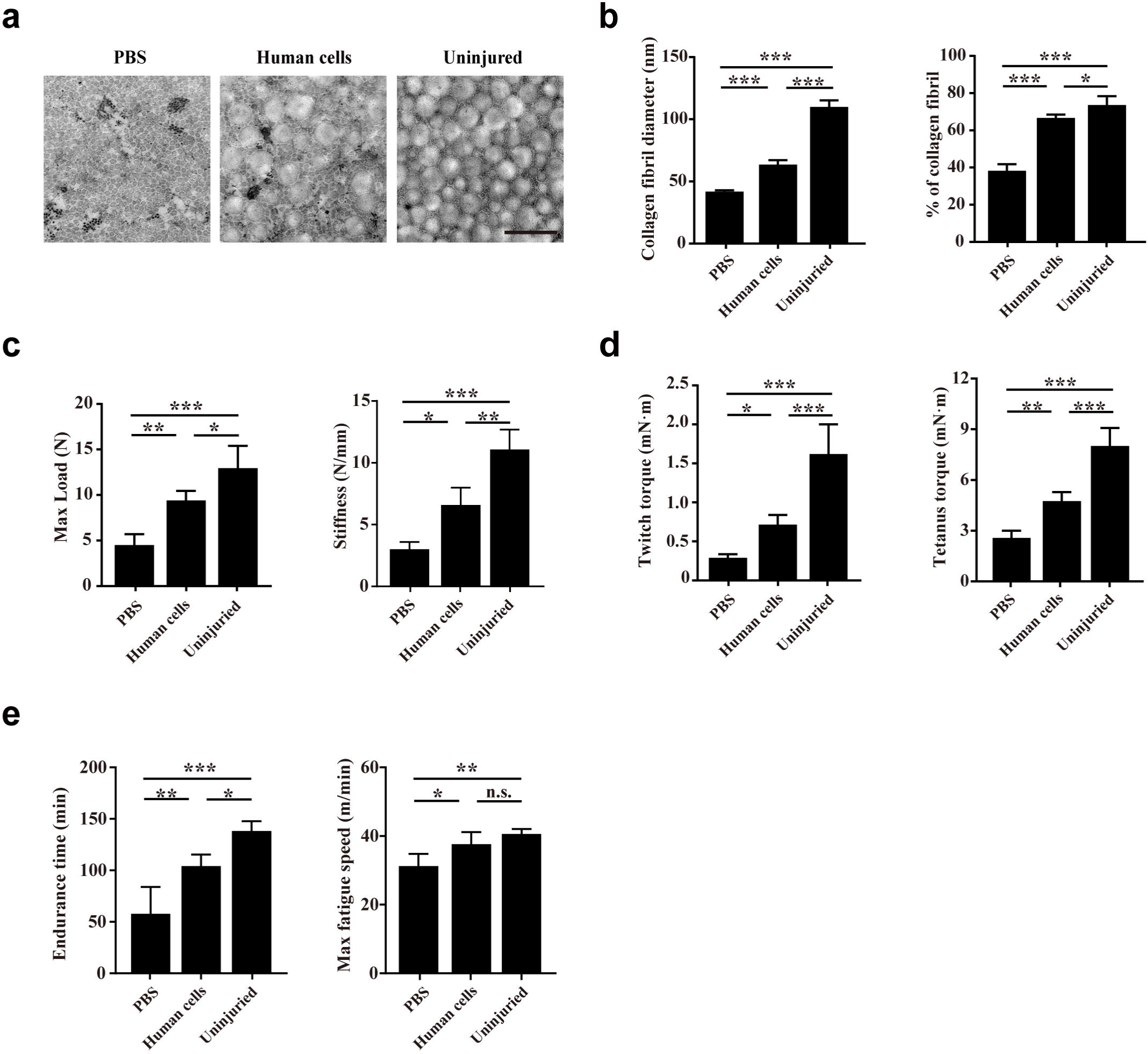
Transplantation of human CD29+/CD56+ myogenic progenitors improves the locomotor function after tendon injury. a. TEM images of collagen fibrils in the injured tendon with PBS injection, injured tendon with human CD29+/CD56+ myogenic progenitors transplantation, and uninjured tendon. Scale bars, 500nm. b. Statistical analysis of the collagen fibril diameter and percentage of collagen fibril area for the injured tendon with PBS injection, injured tendon with human CD29+/CD56+ myogenic progenitors transplantation, and uninjured tendon. Error bars indicated standard deviation (n=5). * indicated p<0.05, *** indicated p<0.001. c. Statistical analysis of the max load and stiffness of the injured tendon with PBS injection, injured tendon with human CD29+/CD56+ myogenic progenitors transplantation, and uninjured tendon. Error bars indicated standard deviation (n=5). * indicated p<0.05, ** indicated p<0.01, *** indicated p<0.001. d. Twitch and tetanus plantarflexion force of the involved limb with PBS injection after tendon injury, human CD29+/CD56+ myogenic progenitors transplantation after tendon injury, and uninjured tendon. Error bars indicated standard deviation (n=5). * indicated p<0.05, ** indicated p<0.01, *** indicated p<0.001. e. The results of treadmill exercise for tendon injured mice with or without human CD29+/CD56+ myogenic progenitors transplantation. The endurance time and max fatigue speed were compared. Error bars indicated standard deviation (n=4). * indicated p<0.05, ** indicated p<0.01, ***indicated p<0.001 and n.s. indicated p>0.05.

Whether the improved tendon regeneration could facilitate the whole organism locomotor function was next investigated. Since the Achilles tendon transmits the plantarflexion force from gastrocnemius muscle to calcaneus, the plantarflexion force of involved leg was also performed two months after tendon injury. Expectedly, transplantation of human CD29+/CD56+ myogenic progenitors for injured tendon also contributed to improving both twitch and tetanus plantarflexion force when compared with PBS injection control group, although the plantarflexion force could still not reach to the level of uninjured leg (Fig. 6d). Consistent with the improved plantarflexion force, the endurance time and max fatigue speed of the mice transplanted with human CD29+/CD56+ myogenic progenitors were better in treadmill test when compared to PBS group (Fig. 6e). Since immunofluorescence staining with human Lamin A/C revealed that only a small number of human CD29+/CD56+ myogenic progenitors engrafted in muscle sporadically (Fig. 5c and d), these results suggest that the transplanted human CD29+/CD56+ myogenic progenitors can improve locomotor function by directly repairing injured tendon.

### TGF**β** signaling pathway contributes to tenogenic differentiation of human CD29+/CD56+ myogenic progenitors

Since the human CD29+/CD56+ myogenic progenitors and murine MuSCs shared the same strain of recipient mice and the same tendon injury while being transplanted, they had the similar microenvironment. Therefore, the distinct differentiation potentials are due to the cell intrinsic differences between species. We further compared the expression profiles of human CD29+/CD56+ myogenic progenitors and murine MuSCs. Interestingly, TGFβ signaling was identified in KEGG enrichment analysis of upregulated genes in human CD29+/CD56+ myogenic progenitors when compared with murine MuSCs (Fig. 7a), indicating that TGFβ signaling could be the key node for maintaining the tendon differentiation potential. Especially, SMAD2 and SMAD3 were identified in upregulated gene set which was enriched in TGFβ signaling pathway (Fig. 7b). Since TGFβ/SMAD2/SMAD3 axis plays a crucial role in tendon development and tenogenic differentiation^22,23,26^, these data further indicated potential crucial role of TGFβ signaling for tenogenic differentiation ability of human CD29+/CD56+ myogenic progenitors. Then TGFβ signaling inhibitor SB-431542 was used to investigate the biological effect of TGFβ signaling during tenogenic induction. The immunofluorescent staining, western blot assay and RT-qPCR results showed SB-431542 significantly suppressed expression level of tendon related genes SCX, TNC, COL I, MKX, and THBS4 (Fig. 7c, d, e). On the contrary, myogenic differentiation ability of human CD29+/CD56+ myogenic progenitors was increased after treatment of SB-431542 during tenogenic induction (Fig. 7f and g). Taken together, these data indicated that TGFβ signaling pathway contributes to tenogenic differentiation of human CD29+/CD56+ myogenic progenitors.

**Figure 7.**
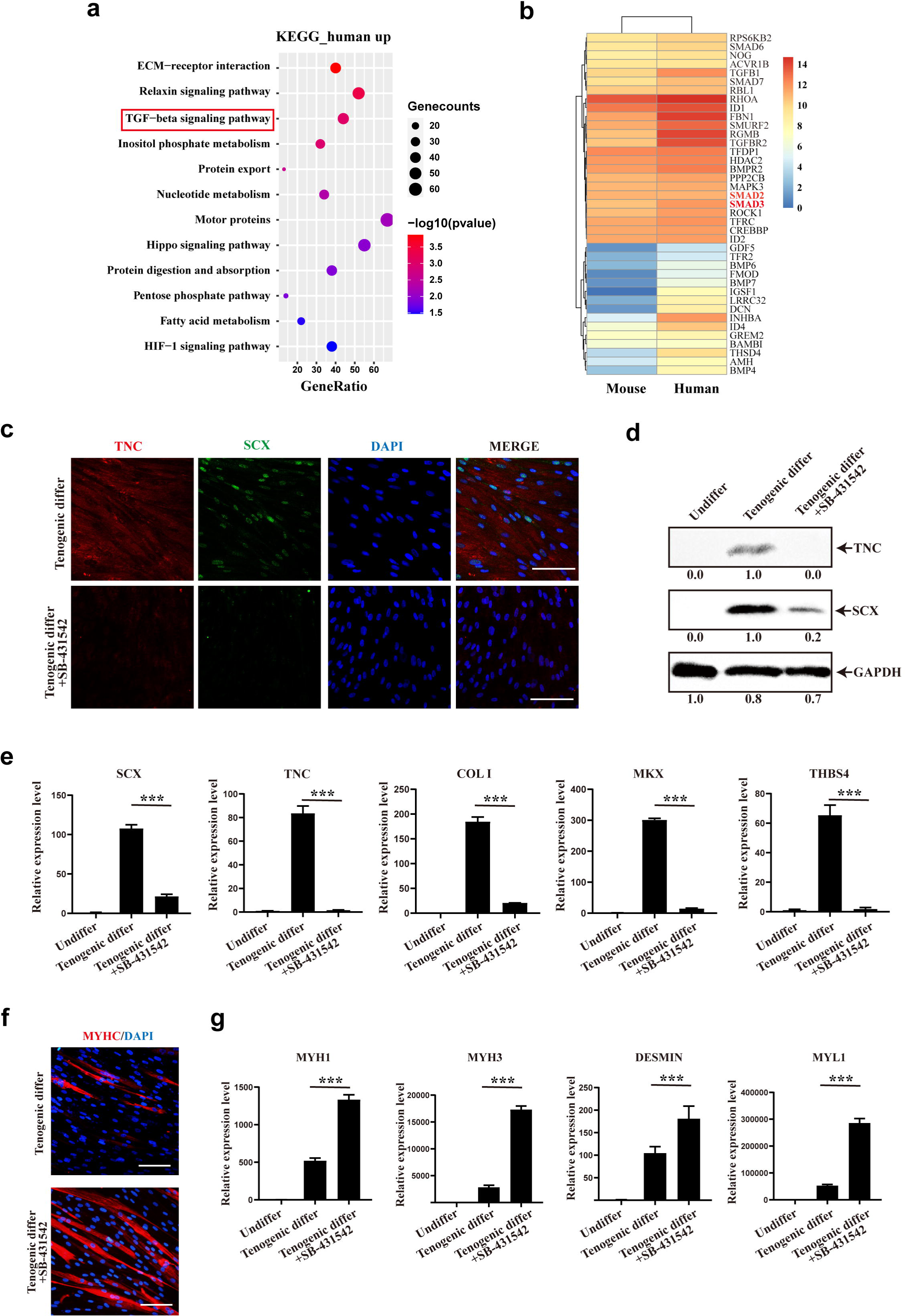
TGFβ signaling pathway contributes to tenogenic differentiation of human CD29+/CD56+ myogenic progenitors. a. Bubble chart of KEGG enrichment analysis of upregulated genes in human CD29+/CD56+ myogenic progenitors when compared with mouse muscle stem cells. b. Heatmap of detailed upregulated genes in human CD29+/CD56+ myogenic progenitors which were enriched in TGFβ signaling pathway. c. Immunofluorescence staining of tendon marker TNC and SCX in human CD29+/CD56+ myogenic progenitors induced for tenogenic differentiation with or without TGFβ signaling pathway inhibitor SB-431542 for 12 days, respectively. Scale bars, 100 µm. d. Protein levels of TNC and SCX. Human CD29+/CD56+ myogenic progenitors were induced towards tenogenic differentiation with or without TGFβ signaling pathway inhibitor SB-431542 for 12 days, respectively. Total protein was extracted from cells before and after differentiation and subjected for TNC and SCX immunoblotting. GAPDH was served as loading control. e. Relative expression levels of tendon related genes. RT-qPCR assays were performed with human CD29+/CD56+ myogenic progenitors upon tenogenic differentiation with or without TGFβ signaling pathway inhibitor SB-431542 for 12 days, respectively. GAPDH was served as reference gene. Error bars indicated standard deviation (n=3). *** indicated p<0.001. f. Immunofluorescence staining of myogenic differentiation marker MyHC in human CD29+/CD56+ myogenic progenitors induced for tenogenic differentiation with or without TGFβ signaling pathway inhibitor SB-431542 for 12 days, respectively. Scale bars, 50 µm. g. Relative expression levels of muscle related genes. RT-qPCR assays were performed with human CD29+/CD56+ myogenic progenitors upon tenogenic differentiation with or without TGFβ signaling pathway inhibitor SB-431542 for 12 days, respectively. GAPDH was served as reference gene. Error bars indicated standard deviation (n=3). *** indicated p<0.001.

## Discussion

Here we show that human CD29+/CD56+ myogenic progenitors have dual differentiation potentials towards muscle and tendon. Transplantation of human CD29+/CD56+ myogenic progenitors contributes to injured tendon repair and thus improves locomotor function. Thus, the human CD29+/CD56+ myogenic progenitors could be served as a new source for tendon regeneration.

Tendon disorders widely occur in people of all ages^22^. It disrupts the stability and mobility of joint, which deeply affects their locomotor function and quality of life. However, the natural healing of injured tendon is very slow due to hypocellularity and hypovascularity of tendon. The biomechanical property and structural integrity could be hardly completely recovered even with surgical treatment^37,38^. It is still a great challenge in clinical work to treat tendon injury.

The relative inefficient outcome of routine therapy for tendon injury sparked the exploration of stem cell treatment. Seed cells with the ability to differentiate into tenocytes and secrete paracrine factors to repair tendon injury are preferred. Thus, tendon derived stem cells, embryonic stem cells, induced pluripotent stem cells and mesenchymal stem cells have been introduced as seed cells to treat tendon injury^39^. However, inadequate sources of tendon derived stem cells, ethical issue and risk of teratoma formation of embryonic stem cells or induced pluripotent stem cell, and heterogeneity of mesenchymal stem cells limit the development of these seed cells for tendon injury treatment. As for muscle progenitors, it is high in proliferation and abundant in sources, and donor site morbidity after muscle harvest was low. Thus, myogenic progenitors might be a promising candidate as seed cells for tendon repair.

Here we also find that the differentiation potential of muscle stem cells is species dependent. Human CD29+/CD56+ myogenic progenitors are bipotential adult stem cells. In contrast, murine muscle stem cells barely have tendon differentiation potential. The species difference might be due to the higher TGFβ signaling level in human CD29+/CD56+ myogenic progenitors. It has been reported that TGFβ/SMAD2/SMAD3 axis plays a crucial role in tendon development and tenogenic differentiation^22,23,26^. SMAD2/3-dependent TGFβ signaling also acts as a crucial molecular brake for myogenesis. It could suppress myogenic regulatory factors Myod1 and Myogenin^40,41^, as well as inhibit myotube fusion and muscle regeneration^42,43^. Thus, the elevated TGFβ signaling may help inhibit the myogenic differentiation ability and stimulate the tenogenic differentiation potential of human CD29+/CD56+ myogenic progenitors under specific microenvironment.

Studies using mouse models contributed to the majority of our knowledge about muscle stem cells and laid the foundation for our understanding of mammalian muscle stem cells. However, human is dramatically different from mouse in many aspects such as size, life span, and manners of motions. It is more demanding for humans to maintain the homeostasis of the locomotion system and the whole organism locomotion ability in much longer life span and bigger body size. Though our knowledge about human muscle regeneration is limited, the current studies have revealed multiple differentiation potentials of Pax7+ human myogenic progenitors in skeletal muscles. CD56+ CD34+ progenitor cells in human skeletal muscle have been reported to have myogenic, osteogenic, and adipogenic activity^14,15^. The CD56+ CD34-progenitor cells in human skeletal muscle have been shown to be free of adipogenic potential^11^. Our results suggest that the CD29+/CD56+ myogenic progenitors in skeletal muscles also have tenogenic differentiation ability besides their myogenic differentiation ability. It seems that there are multiple subpopulations of myogenic progenitors in skeletal muscle. They are all capable of muscle regeneration, while with potentials to regenerate other components of the motion system such as bone, tendon, and adipocytes. This could be an economic method to maintain the functions of the motion system for the longer life span and more complicated motion manner in human beings.

## Methods

### Animals

Animal care and use were in accordance with the guidelines of the animal facility hosted by Shanghai Institute of Biochemistry and Cell Biology, Chinese Academy of Sciences, and the operations were approved by the ethical committee of Shanghai Institute of Biochemistry and Cell Biology. All mice were maintained in specific pathogen-free (SPF) animal facility in individually ventilated cages (IVC) with controlled temperature (22±1℃) and light (12h light/dark cycle). NOD/SCID mice were purchased from Animal Model Research Center of Nanjing University. Pax7CreERT2 and Rosa26-Flox-Stop-Flox-tdTomato mice were purchased from Jackson Laboratory. All experiments were conducted on 3-month-old adult male mice.

### Human samples

Peroneal longus muscle and remanent tendon were obtained from the wastes of patients who underwent full-thickness peroneal longus tendon autograft for knee ligament reconstructive procedures. The study was approved by the ethical committee of Xinhua Hospital Affiliated to Shanghai Jiao Tong University, School of Medicine (Approval No. XHEC-D-2019-043) and written informed consents were obtained from all donors.

### Isolation of human CD29+/CD56+ myogenic progenitors

Human skeletal muscle from peroneal longus were dissected and digested as described previously^6^. Briefly, muscle tissues were cut into small pieces and digested by collagenase II (Worthington biochemical, 700-800U/ml, cat#LS004177) for 60min followed by 30min of digestion with the mixtures of collagenase II and dispase (Life Technologies,11U/ml, cat#17105-041). Digested cells were passed 10 times through a 20-gauge needle. Cell suspension was filtered through a 40µm cell strainer (BD Falcon, cat#352340). Erythrocytes were removed by red blood cell lysis buffer (Thermo Fisher Scientific, cat#00-433-57). The single cell suspension obtained from human muscle was stained with a cocktail containing PE-Cy5 anti-human CD45 (BD Pharmingen, cat#555484), Percp-Cy5.5 anti-human CD31 (BioLegend, cat#303132), AF-488 anti-human CD29 (BioLegend, cat#303016) and PE anti-human CD56 (BioLegend, cat#304606) for 45 min at 4°C. CD45-CD31-CD29+ CD56+ myogenic progenitors were sorted by BD Influx sorter (BD Biosciences).

### Isolation of mouse muscle stem cells

For mouse muscle stem cells isolation, dissected TA muscles were first digested with 10ml muscle digestion buffer (DMEM containing 1% penicillin/streptomycin, 0.125mg/ml Dispase II (Roche, 04942078001), and 10mg/ml Collagenase D (Roche, 11088866001)) for 90 minutes at 37°C. The digestion was stopped by adding 2ml of FBS. The digested cells were filtered through 70μm strainers. Red blood cells were lysed by 7ml RBC lysis buffer (0.802% NH4Cl, 0.084% NaHCO3, 0.037% EDTA in ddH2O, pH7.2-7.4) for 30s, then filter through 40μm strainers. After staining with antibody cocktails (AF700-anti-mouse Sca-1, PerCP/Cy5.5-anti-mouse CD11b, PerCP/Cy5.5-anti-mouse CD31, PerCP/Cy5.5-anti-mouse CD45, FITC anti-mouse CD34, APC-anti-mouse Integrin α7+), the mononuclear cells were subjected for FACS analysis using Influx (BD Biosciences). The population of PI-CD45-CD11b-CD31-Sca1-CD34+ Integrin α7+ cells was collected.

### Primary human tenocytes isolation

Tendon tissues were obtained from the discarded materials of tendon autograft surgery. They were washed with PBS. Epi- and peri-tendon sheath were completely removed. Tenocytes were isolated as described previously^20^. Briefly, the tendons were minced to 1mm^3^ pieces and digested with 3 mg/ml collagenase I (Worthington biochemical, cat# LS004194) in DMEM (Gibco, cat#11965118) at 37℃ for 3hrs with gentle agitation. The digested tissue was filtered through a 40µm cell strainer (BD Falcon, cat#352340) and the isolated cells were plated for subsequent analysis.

### Cell culture and differentiation

Primary human CD29+/CD56+ myogenic progenitors were plated in F10 basal medium (Gibco, cat#11550043) containing 20% FBS (Gibco, cat#10-013-CV), 2.5ng/ml bFGF (R&D, cat#233-FB-025) and 1% Penicillin-Streptomycin (Gibco, cat#15140-122) on collagen-coated dishes. Mouse muscle stem cells were plated in F10 basal medium (Gibco, cat#11550043) containing 20% FBS (Gibco, cat#10-013-CV), 2.5ng/ml bFGF (R&D, cat#233-FB-025) and IL-1α, IL-13, IFN-γ and TNF-α as described previously^36^. DMEM (Gibco, cat#11965118) containing 0.4% Ultroser G (Pall Corporation, cat#15950-17) and 1% Penicillin-Streptomycin were used to differentiate human muscle stem/progenitor cells^5^. DMEM containing 2% horse serum (HyClone, cat#HYCLSH30074.03HI) and 1% Penicillin-Streptomycin were used to differentiate mouse muscle stem cells as described previously^44,45^. DMEM containing 10% FBS, 100ng/ml GDF5 (R&D, cat#8340-G5-050), GDF7 (R&D, cat#8386-G7-050), 0.2mM ascorbic acid (Sigma-Aldrich, cat#A4403) and 1% Penicillin-Streptomycin were used to induce tendon differentiation ^46–48^. Primary tenocytes were cultured in DMEM containing 20% FBS, 2.5ng/ml bFGF (R&D, cat#233-FB-025) and 1% Penicillin-Streptomycin.

### Single-cell RNA sequencing

The single cell suspension of mononuclear cells from human skeletal muscles was firstly prepared. PI (Sangon Biotech, cat#E607328) and Hoechst (Sangon Biotech, cat#A601112) were used to sort live cells by FACS. Then the sored cells were washed twice with PBS containing 0.04% BSA, followed by library preparation with Chromium Single Cell 3’ Reagent Kits (10X genomics, cat# 1000121-1000157). The sequencing was performed on Illumina Novaseq 6000 platform (Illumina).

Single cell RNA-seq data were analyzed by Seurat R (Version 3.2.0) package. Cells with less than 200 genes, more than 6,000 genes detected, and more than 10% mitochondrial genes were excluded. Three samples with a total of 57,193 cells were included for subsequent analysis. Sequencing reads for each gene were normalized to total UMIs in each cell to obtain normalized UMI values by “NormalizeData” function. The “ScaleData” function was used to scale and center expression levels in the data set for dimensional reduction. Total cell clustering was performed by “FindClusters” function at a resolution of 0.1 and dimensionality reduction was performed with “RunUMAP” function^49^.

### Immunofluorescence staining

Cryosections were fixed in 4% formaldehyde for 15min, permeabilized in 0.5% Triton X-100 for 15min, and stained with anti-Pax7 (Developmental Studies Hybridoma Bank), anti-Laminin (Abcam, cat#ab11575), anti-Lamin A/C (Abcam, cat#ab108595; cat#ab190380), anti-TNC (Abcam, cat#ab108930; cat#ab3970), anti-Scx (Abcam, cat#ab58655), anti-Tnmd (Abcam, cat#ab203676), anti-Col I (Abcam, cat#ab260043), anti-Col III (ABclonal, cat#A0817) or anti-MyHC (Millipore, cat#05-716) at 4°C overnight, and incubated with Alexa 488-, Alexa 594 or Alexa 647-labeled anti-mouse or anti-rabbit secondary antibodies (Invitrogen, 1:1000) at room temperature for one hour. The nuclei were stained with 4,6-diamidino-2-phenylindole (DAPI, Vector Laboratories, cat#H-1200). All images were acquired by Leica SP8 confocal microscope (Leica).

### Gene expression analysis

Total RNA was isolated using TRIzol Reagent (Invitrogen, cat#15596-018) according to the manufacturer’s instruction. GAPDH served as internal control. The primers for RT-qPCR are listed as below:

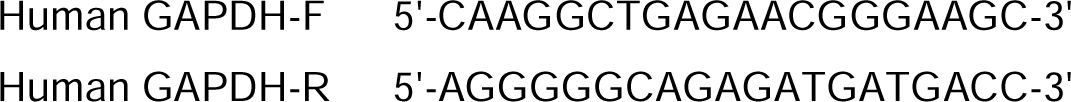
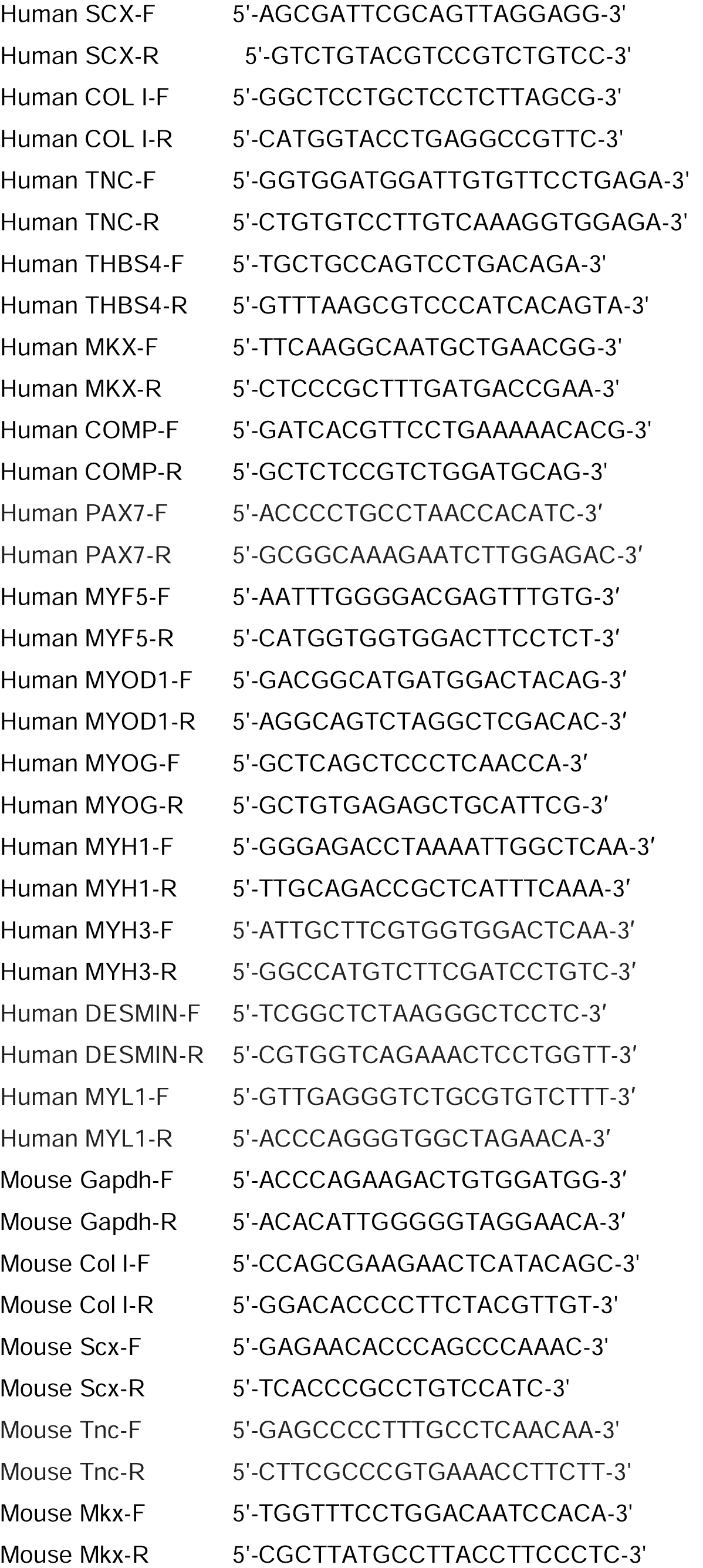
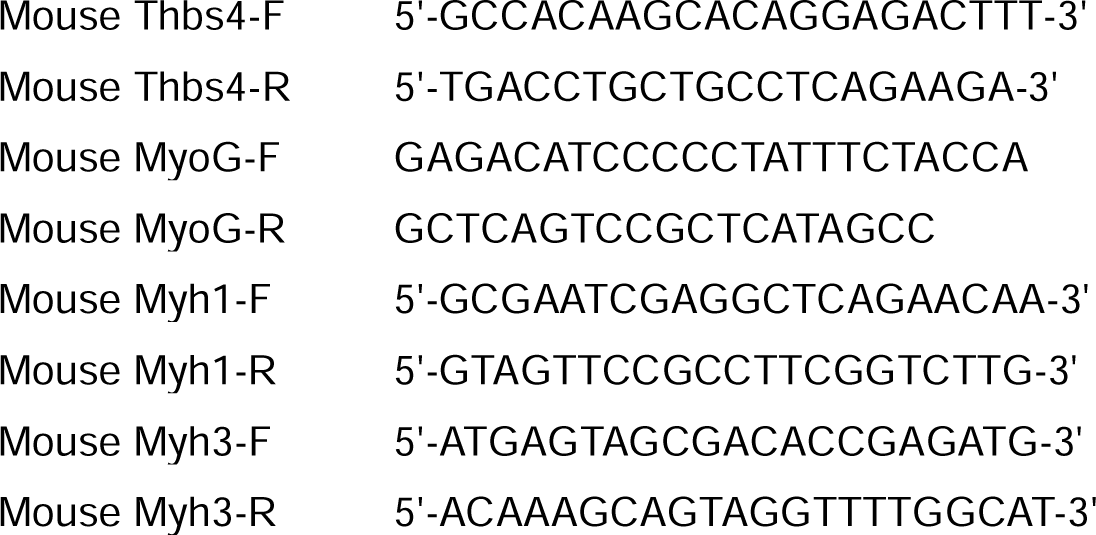

### Cloning assay

Primary human CD29+/CD56+ myogenic progenitors were first sorted in 96 well plates with density of single cell per well. Tenogenic induction or myogenic induction was performed after proliferation for each well. After induction, the immunofluorescence staining of SCX or MyHC was performed in each well. The differentiation efficiency was determined by calculating the ratio of total wells with positive fluorescence signal to total wells with alive cells.

### RNA-sequencing

Total RNA was isolated using TRIzol Reagent (Invitrogen, cat#15596-018) according to the manufacturer’s instruction. mRNA was enriched with magnetic oligo (dT) beads (New England Biolabs, cat#S1419S). The cDNA library was constructed with mean inserts of 200bp with non-stranded library preparation using NEBNext Ultra RNA Library Prep Kit for Illumina (New England Biolabs, cat#E7530L). Sequencing was performed by a paired-end 125 cycles rapid run on the Illumina HiSeq2500. Sequencing data were filtered by SolexaAQ (Q > 20 and length ≥ 25 bp)^50^. The adapter sequences and low quality segments (Phred Quality Score<20) were trimmed using Cutadapt. Paired-end clean reads were then mapped to the reference genome GRCh38.98 using HISAT2. Htseq-count was used to quantify the gene expression value^51^. Read count of each gene was normalized using FPKM (Fragments Per Kilo bases per Million fragments). Differential expression (DE) analysis was performed using DESeq and significant DE genes were defined as those with absolute log2FoldChange>1 and p< 0.05. Heatmap and volcano plot of DE genes was generated by R package Pheatmap (https://cran.r-project.org/web/packages/pheatmap/index.html) and EnhancedVolcano (https://bioconductor.org/packages/release/bioc/vignettes/EnhancedVolcano/inst/doc/EnhancedVolcano.html), respectively. Gene enrichment analysis was conducted by R package topGO with p < 0.05 as the cut-off. The differential expression analysis for RNA-seq data between human CD29+/CD56+ myogenic progenitors and mouse MuSCs was performed according to previous literature^52^.

### Cell transplantation in TA muscle

A single dose of 18 Grey irradiation was administered to the hind legs of the recipient NOD/SCID mice. TA muscle was injured by injecting 15μl of 10μM CTX (Sigma), and 50,000 human CD29+/CD56+ myogenic progenitors or 50,000 murine MuSCs suspended in 10μl PBS were injected intramuscularly to the injury site as described ^5^.

### Tendon injury for lineage tracing

Pax7^CreERT^^2^:Rosa26^tdTomato^ mice were obtained and MuSC was labeled with tdTomato as aforementioned. We used dedicated surgical instruments to imitate a mouse model similar to previously published human surgery^32^. Firstly, distal gastrocnemius tendon was woven with a 6-0 polypropylene non-absorbable suture (PROLENE, cat#EH7242H) and released by microsurgical scissor. After that, a dedicated mini tendon stripper was introduced over the free end of distal medial gastrocnemius tendon. The medial gastrocnemius tendon could be totally removed after advancement of tendon stripper. To further activate MuSCs to guarantee that sufficient amount of activated MuSCs were available around the tendon injury site, 15μl of 10μM CTX cardiotoxin (CTX) was injected at the muscle adjacent to the sites where the tendon was removed to induce more muscle injuries and further activate MuSCs. Then the subcutaneous tissue and skin were sutured.

### Tendon injury and cell transplantation

An approximately 1.5mm long and 0.5mm width transverse incision was performed at 5mm from the calcaneus in Achilles tendon for recipient NOD/SCID mice. 50,000 human CD29+/CD56+ myogenic progenitors or mouse MuSCs resuspended in 20μl hydrogel were injected to the injury site with 28-gauge needles. PBS mixed with 20μl hydrogel were injected as control.

### Biomechanical analysis

Two months after tendon injury, the tendons were harvested to biomechanical analysis using a universal tester (Instron 3345 load system, USA). The grippers were gradually moved apart with the speed of 5 mm/min until the tested tendon was completely ruptured. Then the max load was obtained and documented. The slope of the stress-strain curve was defined as stiffness. All these data were automatically presented in Instron 3345 load system.

### Transmission electron microscope (TEM) examination

After washing the selected target tendon tissues with PBS, tissue fixation was performed for more than 2 hours. Post-fixation was conducted by 1% OsO_4_ (Ted Pella Inc) in 0.1M PB for 1-2 hours. Then dehydration and drying were performed and target tendon samples were subsequently attached to metallic stubs for conductive metal coating. Images were obtained by TEM (HITACHI, SU8100) and at least 1000 collagen fibrils were evaluated for each sample. Density of fibrils was evaluated by the percentage of collagen fibril containing area.

### Treadmill test

Mouse treadmill test was performed in Exer-3/6 treadmill apparatus with electrical stimulus. Treadmill exercise began with a 5 min warm-up, then each mouse was evaluated with specific protocol. For endurance test, mice ran on the treadmill at the constant speed of 22m/min until exhaustion. For exhaustion test, the treadmill speed was increased (2m/min each 3 min) at the beginning speed at 18m/min until exhaustion. Exhaustion was defined as inability to run on the treadmill for longer than10 seconds.

### In vivo muscle force analysis

The 1300A 3-in-1 whole animal system (Aurora Scientific) was used for in vivo muscle force analysis. Mice were first anesthetized and kept warm by heat lamp. The foot was placed on a footplate and kept perpendicular to the tibia. The peroneal branch of the sciatic nerve was stimulated to evaluate the plantarflexion strength. Five repetitive tests were performed for each limb and DMA software (Aurora scientific) was used for results analysis.

### Statistical analysis

At least three biological replicates and technical repeats were performed in each experimental group. All experiments were analyzed and evaluated by investigators in a blinded manner. Error bars indicated standard deviation. Two-tailed unpaired Student’s t-tests were used when variances were similar (tested with F-test) for comparison of two independent groups. One-way ANOVA tests followed by Dunnett’s post-test or Tukey’s post-test were used for multiple comparisons. Shapiro-Wilk tests were performed to determine data normality. Statistical analysis was performed in GraphPad Prism 7 (GraphPad Software, San Diego, USA) or SPSS version 19.0 for Windows (SPSS Inc., Chicago, IL, USA). It was considered significant with P value less than 0.05. Data were presented as mean ± standard unless stated.

### Accession numbers

The complete sequencing data have been uploaded on to Sequence Read Archive database (PRJNA1178160, PRJNA1012476 and PRJNA1012828).

## Supporting information

Shao-Supplemental Figure 1-Fig2 related

Shao-Supplemental Figure 2-Fig4 related

Shao-supplemental Figure 3-Fig5 related

## Acknowledgements

We thank Dr. Dangsheng Li for helpful discussions, the National Protein Science Center (Shanghai) for helps on FACS sorting, the cell biology facility of SIBCB for helps on imaging and FACS analysis. This work was supported by the Strategic Priority Research Program of the Chinese Academy of Science (XDA16020400), National Science Foundation of China (32170804, 81871096, 82372384, 82302657).

## Supplemental Figure legends

**Supplemental Figure 1.**
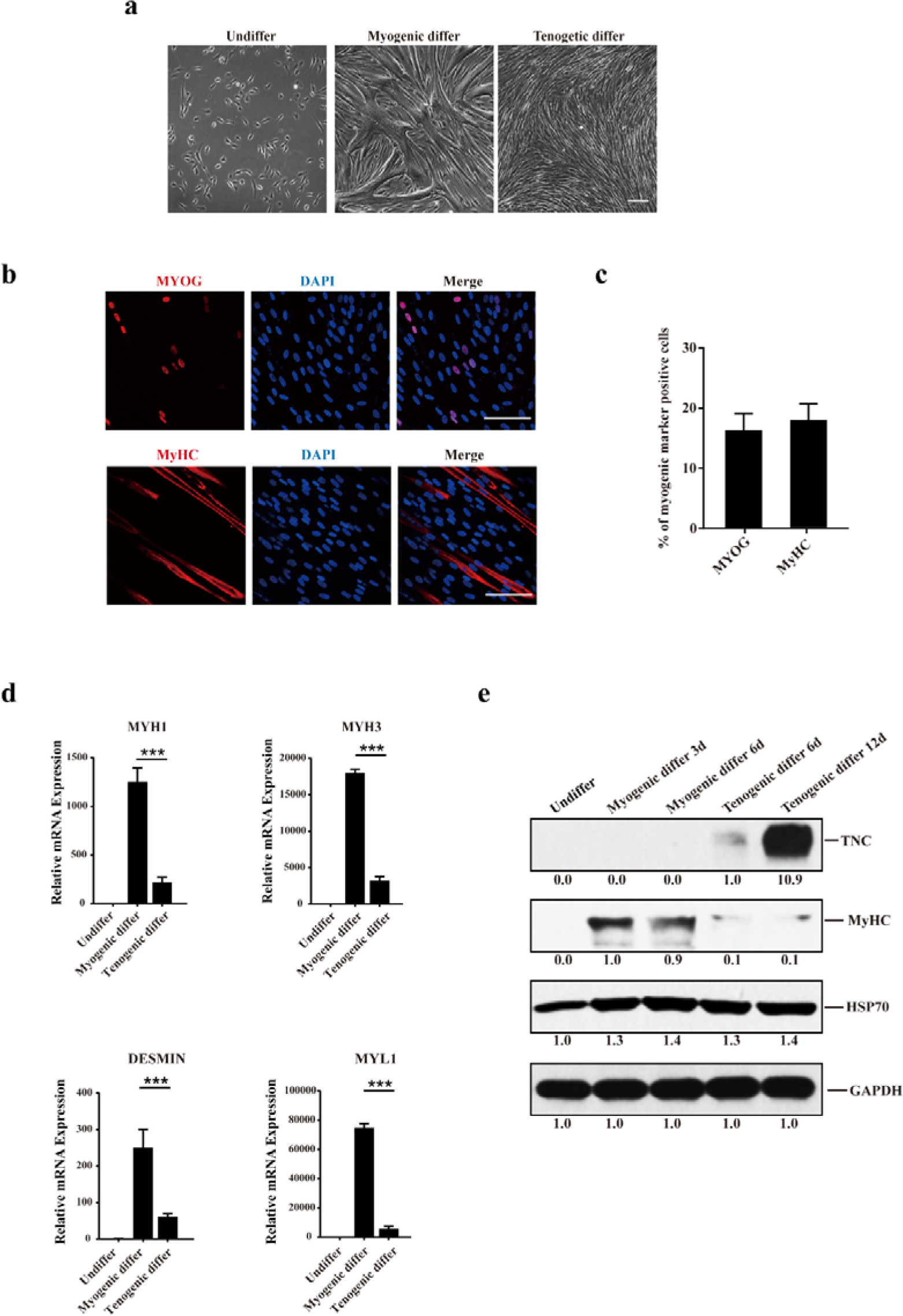
Huan CD29+/CD56+ myogenic progenitors display tenogenic differentiation potential. a. Phase contrast images of human CD29+/CD56+ myogenic progenitors before and after differentiation. Scale bars, 100 µm. b. Immunofluorescence staining of myogenic differentiation marker genes MYOG and MyHC. Human CD29+/CD56+ myogenic progenitors were induced for tenogenic differentiation. The differentiated cells were stained with MYOG and MyHC antibody. Red indicated MYOG or MyHC; DAPI indicated nuclei staining; merge indicated the merged images of red and blue staining. Scale bars, 100µm. c. Statistical analysis of the percentage of human cells expressing MYOG or MyHC under tenogenic differentiation condition. Error bars indicated standard deviation (n=5). d. Expression level of genes marking myogenic differentiation. Human CD29+/CD56+ myogenic progenitors were induced for myogenic and tenogenic differentiation, respectively. The undifferentiated and differentiated cells were subjected for RT-qPCR analysis of MYH1, MYH3, DESMIN, and MYL1. GAPDH was served as reference gene. Error bars indicated standard deviation (n=5). *** indicated p<0.001. e. Protein levels of TNC and MyHC. Human CD29+/CD56+ myogenic progenitors were induced towards myogenic and tenogenic differentiation, respectively. Total protein was extracted from cells before and after differentiation and subjected for TNC and MyHC immunoblotting. HSP70 and GAPDH were served as loading controls.

**Supplemental Figure 2.**
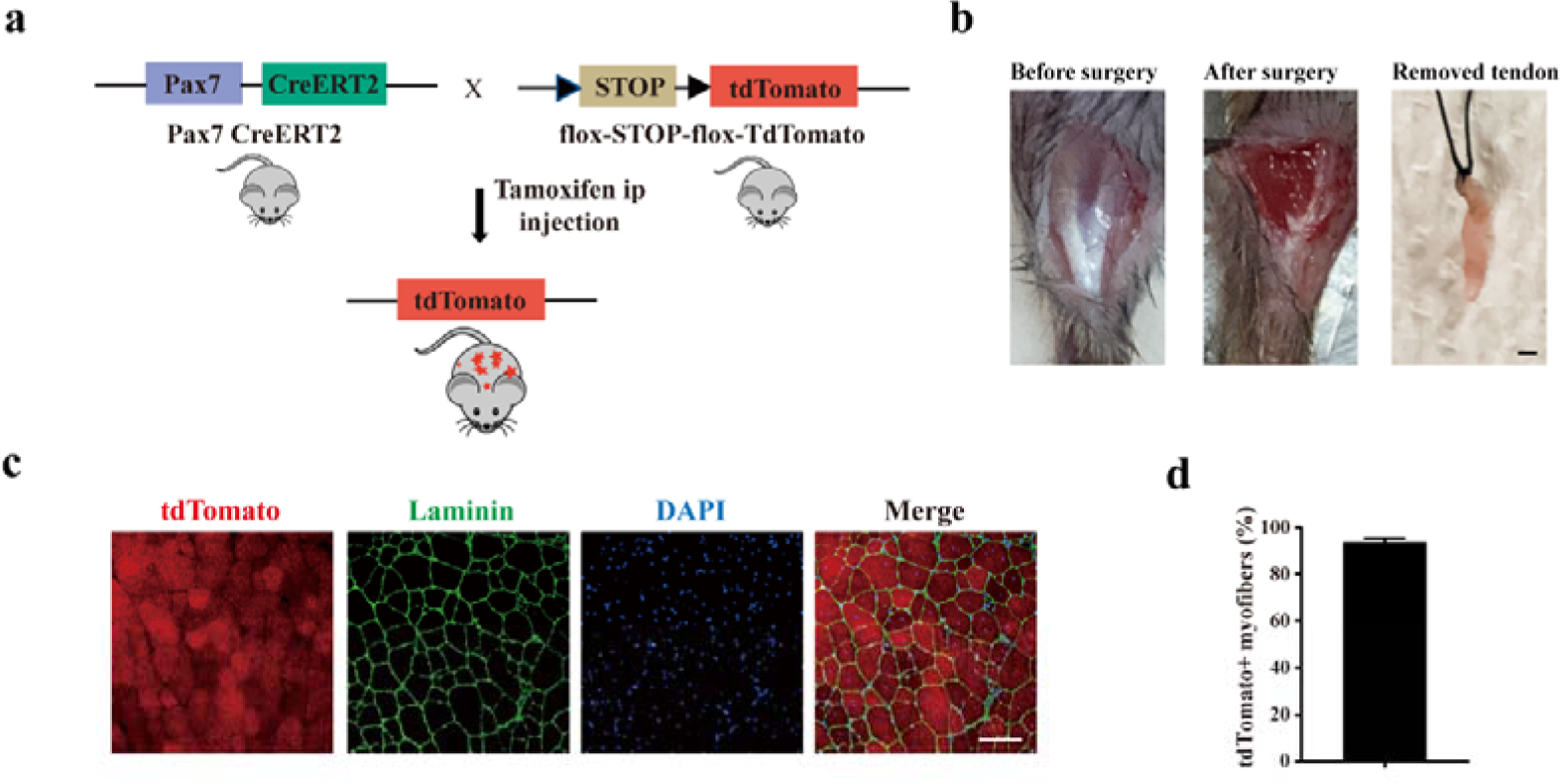
Murine MuSCs display poor tenogenic differentiation ability. a. Schematic illustration of the mouse line for murine MuSCs lineage tracing. b. The surgery in mice mimicking human peroneus longus tendon removal surgery. The medial gastrocnemius tendon was completely removed. Scale bars, 1mm. c. Immunofluorescence images of injured muscle adjacent to the removed tendon. The MuSCs were labeled with tdTomato before injury. The tdTomato positive myofibers after muscle injury were analyzed to evaluate the tracing system. Scale bars, 100μm. d. Statistical analysis of the tdTomato positive myofibers adjacent to the removed tendon. Error bars indicated standard deviation (n=5).

**Supplemental Figure 3.**
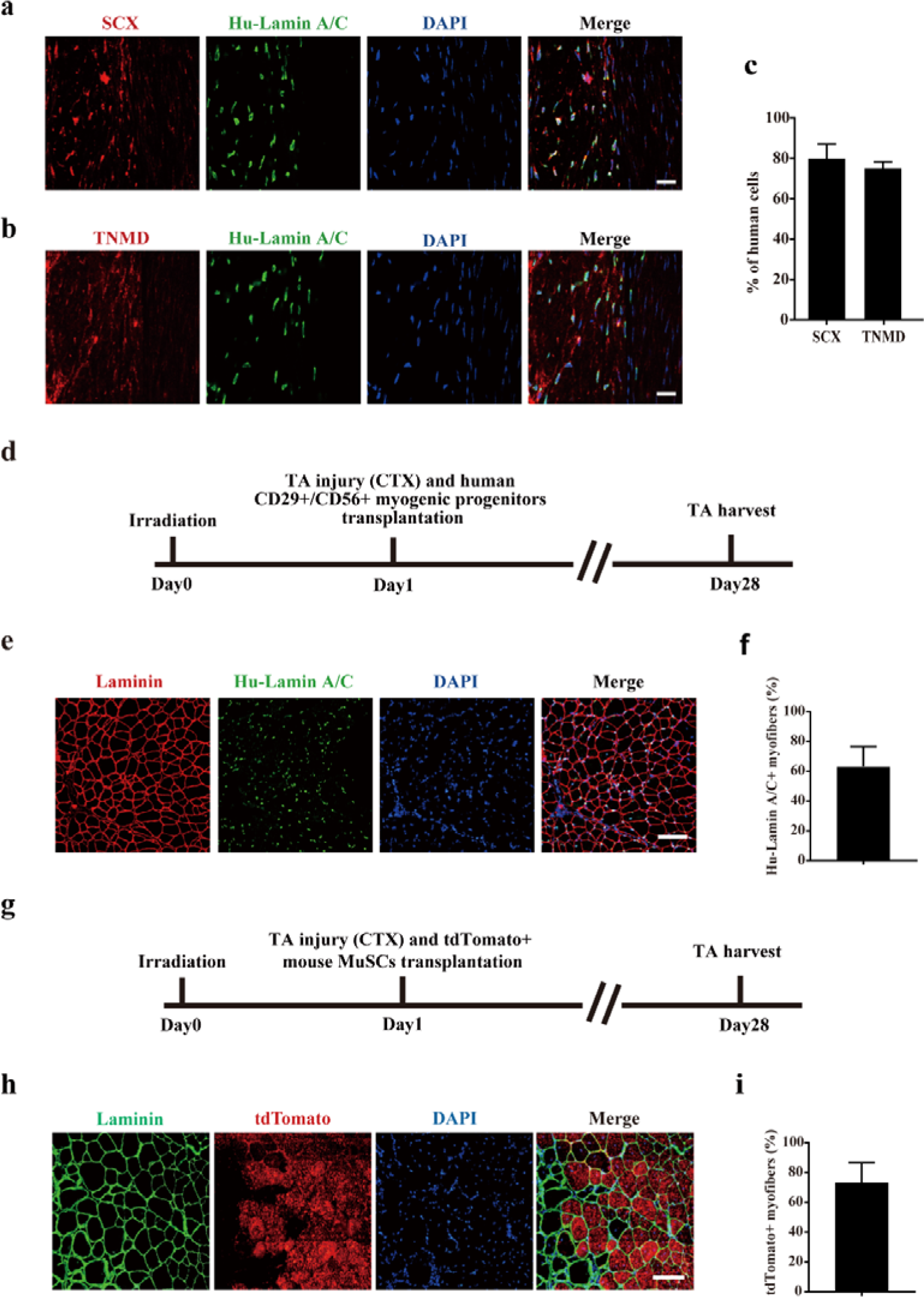
Transplantation of human CD29+/CD56+ myogenic progenitors facilitates tendon regeneration and transplantation of human/murine myogenic progenitors improve muscle regeneration. a. Immunofluorescence staining of SCX and human specific Lamin A/C in injured tendon. Two months after transplantation of human CD29+/CD56+ myogenic progenitors, the injured tendon was stained with SCX and human specific Lamin A/C. DAPI indicated nuclei staining. Merge indicated the merged images of SCX, human specific Lamin A/C, and DAPI. Scale bars, 20 µm. b. Immunofluorescence staining of TNMD and human specific Lamin A/C in injured tendon. Two months after transplantation of human CD29+/CD56+ myogenic progenitors, the injured tendon was stained with TNMD and human specific Lamin A/C. DAPI indicated nuclei staining. Merge indicated the merged images of TNMD, human Lamin A/C, and DAPI. Scale bars, 20 µm. c. Statistical analysis of the percentage of human cells expressing tendon related markers SCX or TNMD after being transplanted to the injured tendon. Error bars indicated standard deviation (n=5). d. Scheme of transplantation of human CD29+/CD56+ myogenic progenitors after muscle injury. TA muscles were first irradiated to kill the local MuSCs in NOD/SCID mice. Then transplantation of human CD29+/CD56+ myogenic progenitors to the irradiated pre-injured recipient mice was performed and TA muscles were harvested after 28 days. e. Immunofluorescence staining of MyHC and human specific Lamin A/C after transplantation of human CD29+/CD56+ myogenic progenitors. Scale bars, 100μm. f. Statistical analysis of the engraftment efficiency of human CD29+/CD56+ myogenic progenitors transplantation in muscle. Error bars indicated standard deviation (n=5). g. Scheme of murine MuSCs transplantation after muscle injury. TA muscles were first irradiated to kill the local MuSCs in NOD/SCID mice. Then transplantation of tdTomato positive MuSCs to the irradiated pre-injured recipient mice was performed and TA muscles were harvested after 28 days. h. Immunofluorescence staining of Laminin and murine MuSCs constitutively expressing tdTomato after murine tdTomato positive MuSCs transplantation. Scale bars, 100μm. i. Statistical analysis of the engraftment efficiency of murine MuSCs transplantation in muscle. Error bars indicated standard deviation (n=5).

## Data availability

The datasets used in this study can be obtained from the corresponding author upon reasonable request.

## Conflict of interest disclosure

The authors declared no conflict of interest.

## Funding

This work was supported by the Strategic Priority Research Program of the Chinese Academy of Science (XDA16020400), National Science Foundation of China (32170804, 81871096, 82372384, 82302657).

## Contributors

XS, XL, HL, HZ, and XF designed the study, collected the experimental data and wrote the original draft. SL and SZ conducted the image analysis, statistical analysis and edited the manuscript. WY and MW designed the study, reviewed and edited the manuscript. JW and PH administrated the project and edited the manuscript.

## Notes

### Competing Interest Statement

The authors have declared no competing interest.

### Summary of Updates

Xingzuan Lin updated their institutional affiliation and added Minhui Wang and Wenjun Yang as authors. Other relevant content has been modified in accordance with the reviewers' comments.

