## Supplementary figures and images for "Human CD29+/CD56+ myogenic progenitors display tenogenic differentiation potential and facilitate tendon regeneration"

### Shao-Supplemental Figure 1-Fig2 related

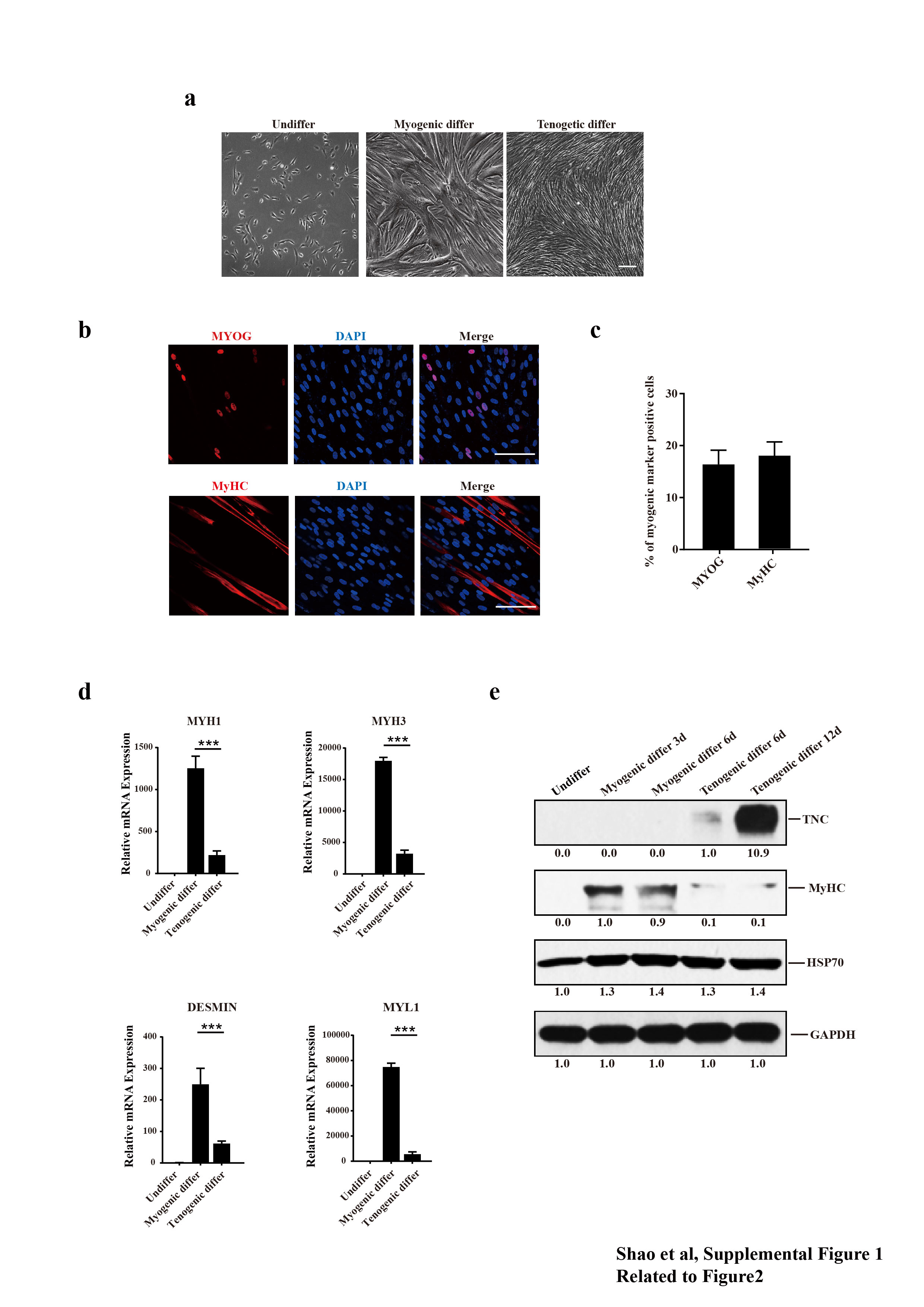

### Shao-Supplemental Figure 2-Fig4 related

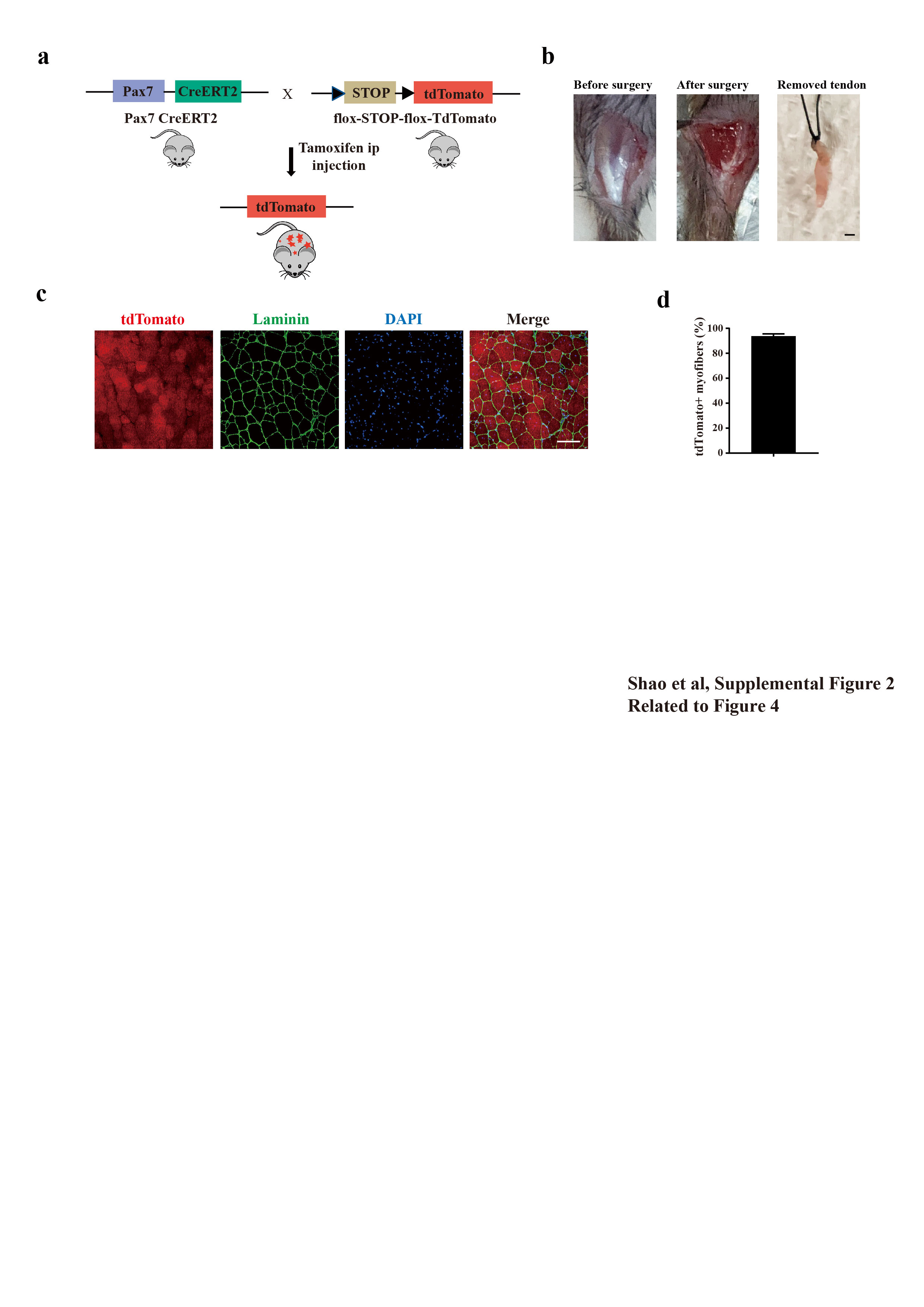

### Shao-supplemental Figure 3-Fig5 related

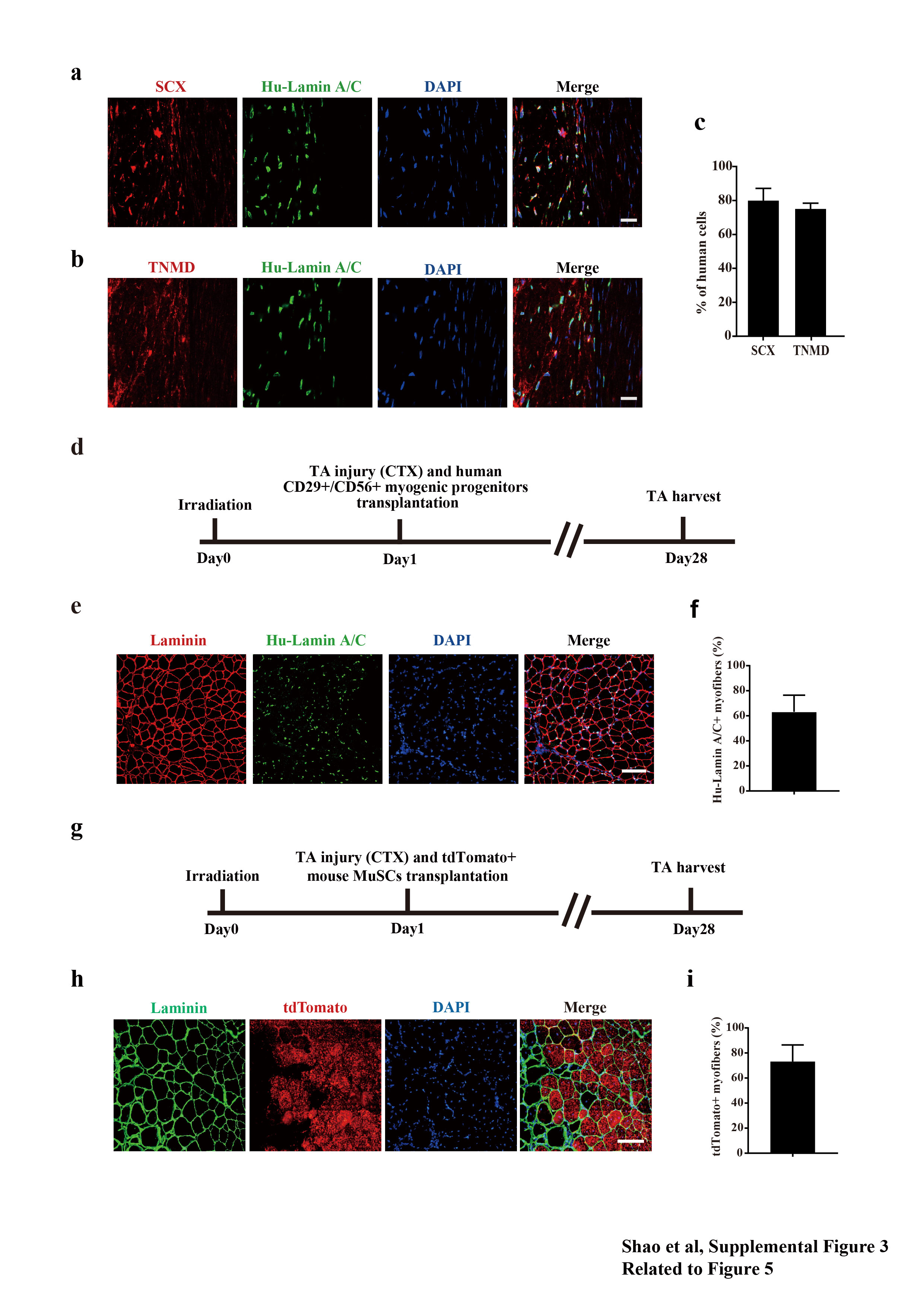
